# Two genetically linked *Arabidopsis* TIR-type NLRs are required for immunity and interact with NLRs encoded in a segmentally duplicated genomic region

**DOI:** 10.1101/2025.04.25.650432

**Authors:** Daniel Lüdke, Qiqi Yan, Max-Emanuel Zirngibl, Charlotte Roth, Melanie Klenke, Annette Gunkel, Marcel Wiermer

## Abstract

Plant nucleotide-binding leucine-rich repeat (NLR) immune receptors detect pathogen-secreted effectors inside host cells and induce a robust immune response, typically involving hypersensitive cell death. *NLR* genes are often genetically linked and can function as pairs or within larger NLR networks. We previously showed that the truncated Toll/Interleukin-1 Receptor (TIR)-type NLR TIR-NB13 (TN13) is required for resistance of *Arabidopsis* to *Pseudomonas syringae* pv. *tomato* (*Pst*) DC3000 lacking the type-III effector proteins AvrPto and AvrPtoB. *TN13* is genetically linked to a full length *TIR-NB-LRR* (*TNL*) gene on chromosome 3. Here, we show that TN13, and its genetically linked TNL both localize to the ER membrane via N-terminal transmembrane domains, are required for resistance to *Pst* DC3000 (ΔAvrPto/AvrPtoB) and interact with each other in transient expression assays in *Nicotiana benthamiana*. In contrast to TN13, the full length TNL, which we named TN13-INTERACTING TNL1 (TNT1), induces an autoactive cell death response when expressed in *N. benthamiana* that depends on an atypical MHV motif in its NB-ARC domain, as well as the EDS1/SAG101/NRG1 module. TN13 and TNT1 furthermore interact with phylogenetically related NLRs encoded by a segmentally duplicated region on chromosome 5. Our data suggest that both TN13 and the genetically linked TNT1 could be part of a larger TIR-type NLR immune regulatory network, in which TNT1 contributes to basal immunity and might function as an autoactive death switch to induce cell death upon pathogen detection.

**SIGNIFICANCE STATEMENT:** The ER membrane localized truncated TIR-NLR TN13 and the genomically linked full length TNL TNT1 are required for plant disease resistance and form heteromeric associations with phylogenetically related NLRs, encoded by a segmentally duplicated chromosomal region.

## INTRODUCTION

Plants evolved a multilayered, cell autonomous immune system to efficiently detect and respond to pathogen threats (Dangl *et al*., 2013). Nucleotide-binding and leucine-rich repeat (NLR) proteins function as intracellular immune receptors to sense pathogen-secreted effector molecules that contribute to pathogen virulence, if they remain undetected (Kourelis and van der Hoorn, 2018; Wu and Derevnina, 2023). Effector recognition by NLRs initiates a robust defense response often including a locally restricted programmed cell death, known as the hypersensitive response (HR), to restrict the growth of biotrophic pathogens that require living host tissues (Dangl *et al*., 2013).

NLRs have a modular, multi-domain architecture with a central NB domain that contains Apaf-1/R-protein/CED4 (ARC1/2) subunits (NB-ARC), a C-terminal LRR-domain and either a Toll/Interleukin-1 receptor (TIR) domain or a coiled-coil (CC) domain at the N-terminus. Thus, NLRs can phylogenetically be divided into distinct classes, referred to as TIR-type NLRs (TNLs) and CC-type NLRs (CNLs; Shao *et al*., 2016; Kourelis *et al*., 2021; Contreras *et al*., 2023).

Plant NLRs belong to the family of signal transduction ATPases with numerous domains (STAND) and are proposed to function as ATP-hydrolyzing molecular switches (Leipe *et al*., 2004; Takken *et al*., 2006; Danot *et al*., 2009). In the absence of a cognate effector molecule or an activating trigger, NLRs are in a resting state and bind ADP with their NB-ARC domain (Williams *et al*., 2011; Wang, Wang, *et al*., 2019; Selvaraj *et al*., 2024). An effector triggered switch to the active state requires the substitution of ADP by ATP (Zhou *et al*., 2015; Wang, Hu, *et al*., 2019; Selvaraj *et al*., 2024). The ADP-ATP exchange and the stabilization of ATP relies on a functional phosphate-binding loop (P-loop) within the NB domain (Wang, Hu, *et al*., 2019). In addition to the P-loop, the histidine of the conserved MHD (Met-His-Asp) motif within the ARC2 domain also interacts with the ADP/ATP β-phosphate (Reubold *et al*., 2011). An amino acid substitution from MHD to MHV can render plant NLRs autoactive, presumably by shifting the binding preference from ADP towards ATP (Bendahmane *et al*., 2002; Tameling *et al*., 2006; Williams *et al*., 2011). The nucleotide-bound state also correlates with intra-molecular interactions between different NLR domains (Ve *et al*., 2015; Bentham *et al*., 2018). The LRR domain directly interacts with parts of the NB-ARC domain to maintain the NLR in a repressed state (Slootweg *et al*., 2013; Schreiber *et al*., 2016). Direct interactions with effectors or sensing of effector-induced host protein modifications by LRR domains can release intra-molecular auto-inhibitory effects for ATP binding and NLR activation (Slootweg *et al*., 2013; Ntoukakis *et al*., 2013).

Plant NLRs rarely function as singletons, which are capable of both effector detection and cell death induction, but rather form NLR pairs or complex networks which separates effector detection and downstream signaling and immunity (Lüdke *et al*., 2022; Contreras *et al*., 2023). Accordingly, sensor NLRs perceive effectors, and signal through helper NLRs to initiate immune signaling (Qi *et al*., 2018; Wu *et al*., 2017; Castel *et al*., 2019; Lüdke *et al*., 2022; Contreras *et al*., 2023). All currently identified full length TIR-type NLRs act as sensors that either signal through ADR1 (ACTIVATED DISEASE RESISTANCE1) or NRG1 (N REQUIREMENT GENE1) as helper NLRs for induction of immunity or HR (Wu *et al*., 2019; Lapin *et al*., 2019; Saile *et al*., 2020). Inhibitory inter-molecular interactions between sensor NLR pairs are an additional regulatory layer to prevent signaling in the absence of a pathogen (Césari *et al*., 2014; Huh *et al*., 2017). For instance, effector perception by the inhibitory *Arabidopsis* TNL sensor RESISTANCE TO RALSTONIA SOLANACEARUM1 (RRS1-R) leads to dissociation from the TNL partner RESISTANCE TO PSEUDOMONAS SYRINGAE4 (RPS4), which acts as an executer to induce an immune response that depends on the helper NLR NRG1 (Sarris *et al*., 2015; Le Roux *et al*., 2016; Huh *et al*., 2017; Ma *et al*., 2018; Castel *et al*., 2019). The helper NLR classes of NRG1s and ADR1s form a subgroup of NLRs, in which the N-terminal CC domain resembles the CC motif of the resistance protein RESISTANCE TO POWDERY MILDEW8 (RPW8; Peart *et al*., 2005; Bonardi *et al*., 2011; Dong *et al*., 2016). While some *Arabidopsis* NLRs require both ADR1 and NRG1, others require either NRG1 or ADR1 for immune signaling (Castel *et al*., 2019; Wu *et al*., 2019). Besides helper NLRs, the defense regulator ENHANCED DISEASE SUSEPTIBILITY1 (EDS1) is essential for TIR-type NLR mediated immunity and is also required for signaling of some CNLs (Wiermer *et al*., 2005; Xiao *et al*., 2005; Wagner *et al*., 2013). The EDS1-related proteins PHYTOALEXIN DEFICIENT4 (PAD4) and SENESCENCE ASSOCIATED GENE101 (SAG101) function together with EDS1 and form distinct hetero complexes (Feys *et al*., 2005; Huang *et al*., 2022; Feehan *et al*., 2023). Whereas signaling via an EDS1-PAD4-ADR1 module restricts bacterial growth through transcriptional reinforcement, EDS1-SAG101-NRG1 were proposed to signal mainly for HR induction (Bonardi *et al*., 2011; Dong *et al*., 2016; Cui *et al*., 2018; Castel *et al*., 2019; Lapin *et al*., 2019; Saile *et al*., 2020; Feehan *et al*., 2023). Following effector detection by TIR-type NLR sensors, NRG1 and ADR1 helper NLRs oligomerize and localize to the plasma membrane where they mediate Ca^2+^ influx to induce immune signaling (Jacob *et al*., 2021; Feehan *et al*., 2023). Activated NRG1 was also shown to target other organellar membranes, such as the chloroplast (Ibrahim *et al*., 2024).

TIR-type NLR sensors function as NAD^+^/NADP^+^ degrading enzymes and produce cyclic ADP-ribose variants (v-cADPR) that are required for cell death induction (Horsefield *et al*., 2019; Jia *et al*., 2022). While ADP-ribosylated ATP (ADPr-ATP) and ADPr-ADPR (di-ADPR) binding to EDS1-SAG101 dimers promotes NRG1 association and oligomerization (Jia *et al*., 2022), 2′-(5′-phosphoribosyl)-5′-adenosine mono-/di-phosphate (pRib-AMP and pRib-ADP) bind to EDS1-PAD4 dimers leading to ADR1 association and oligomerization (Huang *et al*., 2022). In addition, it was recently shown that the 2′,3′-cAMP/cGMP synthetase activity of TIR domain proteins such as the TIR-only NLR RESPONSE TO THE BACTERIAL TYPE III EFFECTOR PROTEIN HOPBA1 (RBA1) also contributes to cell death induction (Yu *et al*., 2022). RBA1 is required for immunity to *Pseudomonas syringae* expressing the effector protein HopBA1, and induces an HR cell death response when expressed transiently in *Nicotiana benthamiana* (Nishimura *et al*., 2017). Other truncated NLRs encoded by the *Arabidopsis* genome include TIR-NB (TN) and TIR-X (TX) proteins that lack NB and/or LRR domains but might contain additional (X) domains (Meyers *et al*., 2002; Nandety *et al*., 2013; Van de Weyer *et al*., 2019). A functional role of truncated NLRs in immunity has been described for TN2, which is required for defense signaling in the exo*70b1-3* auto-immune mutant, suggesting that TN2 monitors the integrity of EXOCYST SUBUNIT EXO70 FAMILY PROTEIN B1 (EXO70B1), involved in vesicle exocytosis (Zhao *et al*., 2015). In addition, TN2 monitors the cellular homeostasis of the E3 ligase SENESCENCE-ASSOCIATED E3 UBIQUITIN LIGASE1 (SAUL1) together with the TN-type NLR CHILLING SENSITIVE1 (CHS1/TN1), both of which depend on the TNL SUPPRESSOR OF CHS1 (SOC3; Zhang *et al*., 2017; Tong *et al*., 2017; Liang *et al*., 2019). Significantly, *SOC3* forms a genomic cluster with *CHS1* and *TN2*, a chromosomal arrangement commonly observed among NLRs that function cooperatively (Meyers *et al*., 2003; van Wersch and Li, 2019).

We previously showed that the ER-localized truncated NLR TIR-NB13 (TN13) is required for immunity of *Arabidopsis* to mildly virulent *Pseudomonas syringae* pv. *tomato* (*Pst*) DC3000 (Roth *et al*., 2017). The gene encoding TN13 (*AT3G04210*) is linked genetically in a head-to-tail orientation to a predicted full length *TNL* gene (*AT3G04220*).

Here, we report that AT3G04220, which we named TN13-INTERACTING TNL1 (TNT1), interacts and co-localizes with TN13 at the ER membrane via its N-terminal transmembrane domain, and is also required for immunity to mildly virulent *Pst* DC3000. Interestingly, TNT1 contains an atypical MHV motif in its wild-type amino acid sequence, which is associated with autoactivity and appears not to be present in other *Arabidopsis* NLRs. We show that this motif is required for the induction of HR when expressed transiently in *N. benthamiana*. The cell death response further depends on a predicted NAD^+^/NADP^+^ degrading catalytic glutamate residue present in the TIR domain, and the *NbEDS1a*/*NbSAG101b*/*NbNRG1*, but not on the *NbPAD4/NbADR1* module. The genomic region containing *TN13* and *TNT1* on chromosome 3 was segmentally duplicated in the *Arabidopsis* genome and contains an expanded number of *TNL* genes on chromosome 5. TN13 and TNT1 interact with each other and with phylogenetically closely related NLRs of the duplicated region, suggesting the potential formation of a functional NLR network regulating plant immunity. We propose that TNT1 acts as an autoactive death switch that is regulated by interactions with TN/TNL proteins. Effector detection could activate immune signaling and cell death induction by TNT1 at the cytoplasmic side of the ER membrane.

## RESULTS

### *TN13* and *TNT1* mutants show enhanced susceptibility to *Pst* DC3000 (ΔAvrPto/AvrPtoB)

We previously identified the truncated NLR protein TIR-NB13 (TN13) as a component of immunity against mildly virulent *Pseudomonas syringae* pv. *tomato* (*Pst*) DC3000 lacking the type-III effector proteins AvrPto/PtoB (Roth *et al*., 2017). On *Arabidopsis* chromosome 3, *TN13* is genetically linked in a head-to-tail orientation to a predicted full length *TNL* gene (AT3G04220, Figure 1A) which we named *TN13-INTERACTING TNL1* (*TNT1*). We hypothesized that the genetic linkage suggests cooperative functions in immunity (van Wersch and Li, 2019). We therefore investigated the requirement of *TNT1* in immunity by inoculating homozygous T-DNA insertion mutants with *Pst* DC3000 (ΔAvrPto/AvrPtoB) (Figures 1B, S1). The *tnt1* mutant line showed enhanced susceptibility towards *Pst* DC3000 (ΔAvrPto/AvrPtoB) to an extent that is comparable to the *tn13* mutant line (Figure 1). Consistent with cooperative functions in plant defense against *Pst* DC3000, a *tn13 tnt1* deletion mutant that was generated by CRISPR/Cas9 genome editing, shows susceptibility that is comparable to the *tn13* and *tnt1* single muntants (Figure 1B, S1). The Col *eds1-2* mutant was used as a hyper-susceptible control, whereas the auto-immune mutant *snc1* (*suppressor of npr1-1, constitutive 1*) served as a control for enhanced resistance (Figure 1B). The enhanced susceptibility of the *tn13* and *tnt1* mutant lines was complemented by stable expression of the respective wild-type proteins with a C-terminal mYFP tag, under control of either the native (*Np*) or the constitutively active *35S* promoter, respepctively (Figure S2). Stable transgenic expression of TN13-mYFP by the *35S* promoter resulted in a stunted phenotype commonly associated with activated immunity (Figure S2A). The *tn13* and *tnt1* mutant lines and the *tn13 tnt1* double mutant appeared wild-type-like and did not display signs of autoimmunity (Figure 1, S2A). We conclude that the truncated TIR-type NLR *TN13*, and the genomically linked full length NLR *TNT1*, are both required for immunity against mildly virulent *Pseudomonas* bacteria.

**Figure 1.**
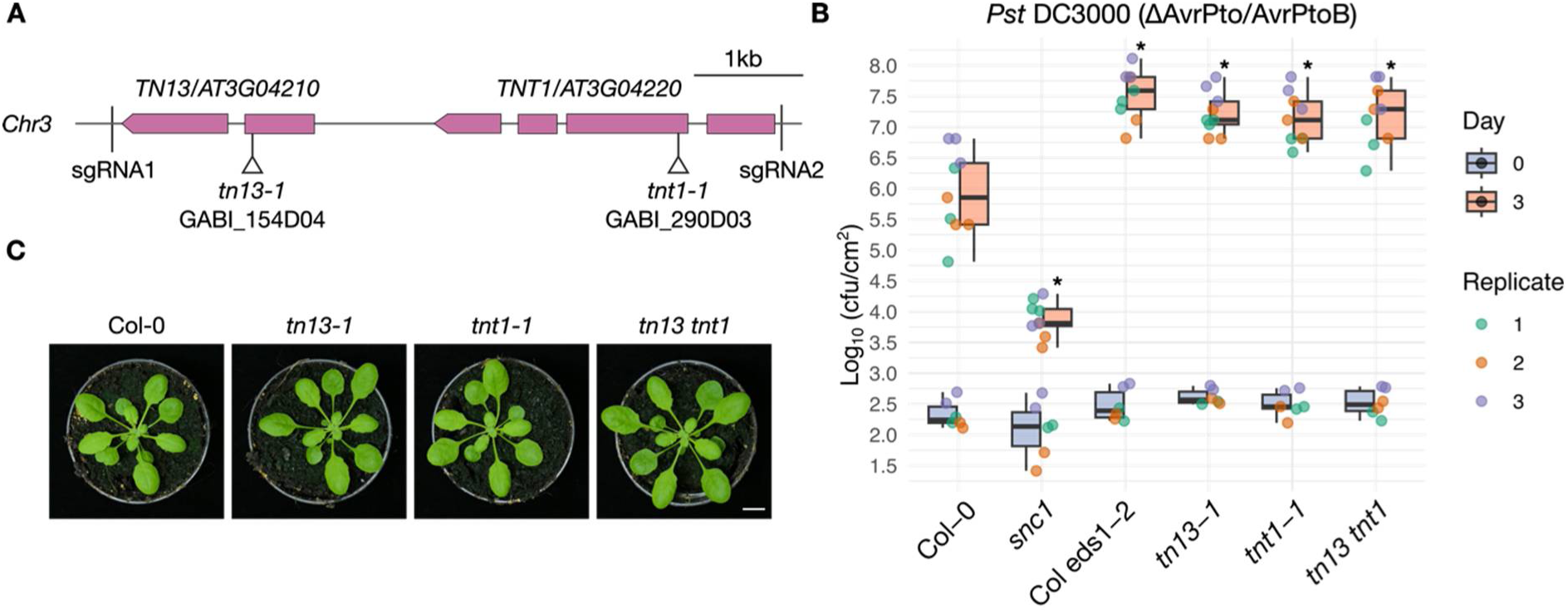
*TNT1* and *TN13* mutants show enhanced susceptibility to *Pseudomonas syringae* pv. *tomato* (*Pst*) DC3000 (ΔAvrPto/AvrPtoB). (A) Gene structures of genetically linked *TN13* and *TNT1* drawn to scale. Exons are represented as boxes and introns as solid lines. The positions of the T-DNA insertions in mutant lines used in infection studies are marked by triangles below the gene structures. The position of single guide RNAs (sgRNA) used to generate the *tn13 tnt1* double mutant are indicated. (B) Plants of the indicated genotypes were infiltrated with a *Pst* DC3000 (ΔAvrPto/AvrPtoB) suspension of 1 x 10^5^ cfu ml^-1^. Colony-forming units (cfu) were quantified 1 hour (day 0) and three days after infiltration (day 3). Data is presented as boxplots, datapoints are colored based on replicates. Asterisks indicate statistically significant differences to Col-0 (Kurskal-Wallis test; Pairwise Wilcoxon tests with Benjamini-Hochberg correction, *P* < 0.05). (C) Representative images of four-week-old plants of the indicated genotypes grown in parallel on soil under short day (SD) conditions. Scale bar = 1 cm.

### TNT1 induced cell death depends on *EDS1*, *SAG101b* and *NRG1* in *Nicotiana benthamiana*

Based on domain predictions, TNT1 contains canonical TIR-type NLR domains (TIR, NB-ARC and LRR; Figure S3). As observed for TN13 (Roth *et al*., 2017), TNT1 also contains a predicted N-terminal hydrophobic transmembrane (TM) domain, which is predicted to have an alpha helical structure (Figures S3B). We previously reported that TN13 localizes to membranes of the endoplasmic reticulum (ER) and the nuclear envelope after transient expression in *N. benthamiana* (Roth *et al*., 2017). To investigate the subcellular localization of TNT1, we performed transient expression assays of mYFP-tagged TNT1 in *N. benthamiana*. Two days after *Agrobacterium-* infiltration for expression of *35S* promoter or *Np*-driven TNT1-mYFP, we noticed a cell death response, resulting in tissue collapse five days after infiltration (Figure 2A). Cell death was also observed upon transient expression of a C-terminal 3xHA-StrepII-tagged version of TNT1, whereas transient expression of TN13-mYFP did not induce cell death (Figures 2A, S4). To further investigate whether the cell death response depends on EDS1, as previously reported for TIR-type NLRs, we transiently expressed TNT1-mYFP in *Nbeds1a-1* mutant plants (Ordon *et al*., 2017). No cell death could be observed five days after *Agrobacterium*-infiltration, although the full-length fusion protein accumulated (Figures 2B, S4). We utilized *N. benthamiana* single, double and triple mutant combinations of the EDS1 interaction partners PAD4 and SAG101, and the helper NLRs *NRG1* and *ADR1* to elucidate their requirements for cell death induction upon TNT1-mYFP expression. Since an *adr1* mutant was not available at the time, we complemented the *adr1 nrg1* double mutant by co-expressing an untagged version of *NbNRG1* (Lapin *et al*., 2019). HR cell death upon TNT1-mYFP expression fully depends on *NbSAG101b* and *NbNRG1*, whereas *NbSAG101a*, *NbPAD4* and *NbADR1* are dispensable (Figures 2B, S4). The transient expression assays indicate that TNT1 autoactively signals for cell death through the EDS1-SAG101-NRG1 module in *N. benthamiana*.

**Figure 2.**
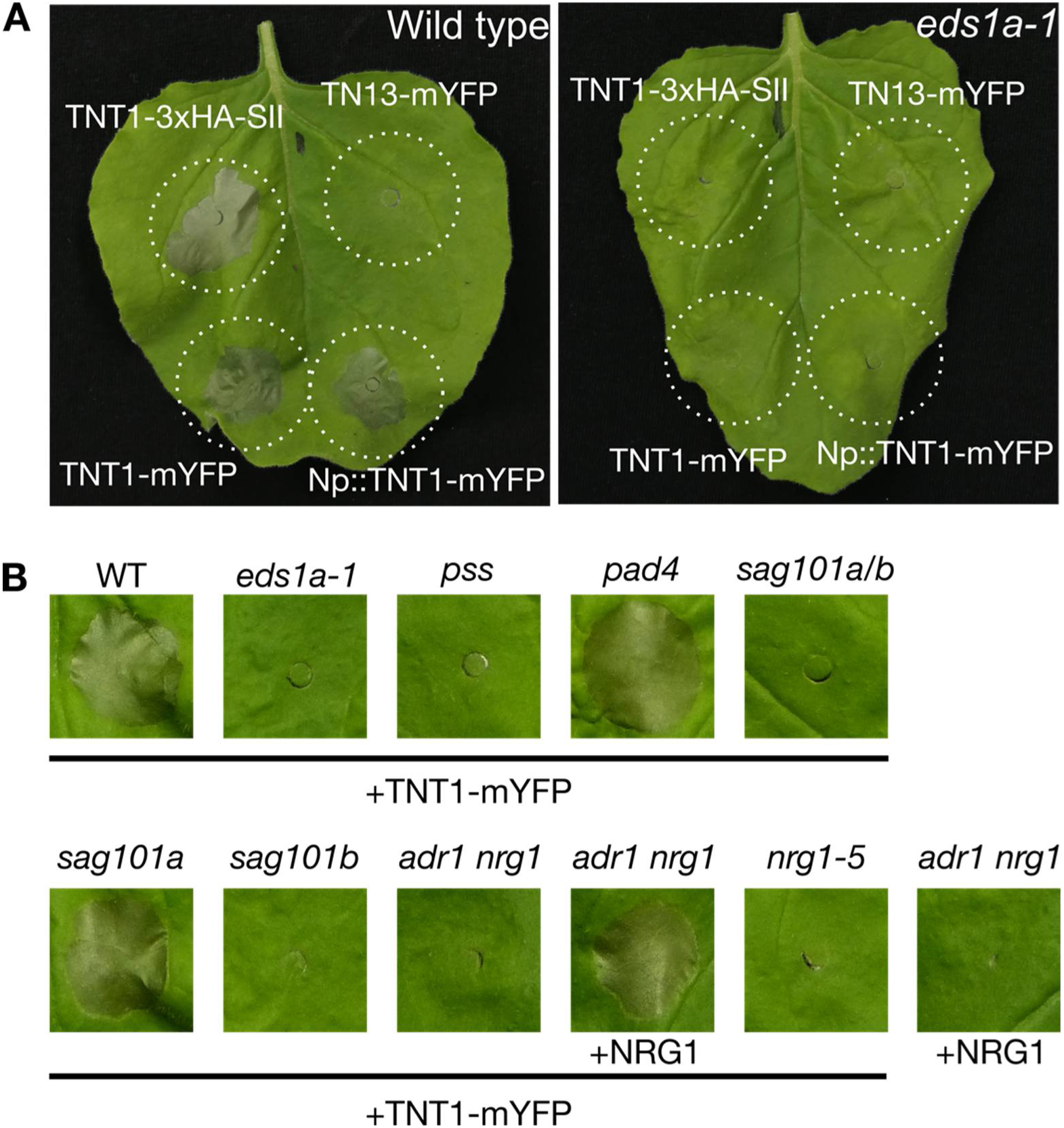
Transient expression of TNT1 in *N. benthamiana* induces a cell death response that depends on *NbEDS1*, *NbSAG101b* and *NbNRG1*. (A) *N. benthamiana* plants of the indicated genotypes were photographed 5d after infiltration with *Agrobacteria* for transient expression of mYFP or 3xHA-StrepII (3xHA-SII) tagged versions of TNT1 or TN13, under control of the *35S* or native (*Np*) promoter. (B) *N. benthamiana* plants of the indicated genotypes were infiltrated with *Agrobacteria* for expression of TNT1-mYFP under control of the *35S* promoter. The *adr1 nrg1* double mutant was complemented with *NRG1* via transient co-expression. *pss*: *pad4 sag101a sag101b*. Cell death was photographed 5d after *Agrobacterium* infiltration.

### TNT1 co-localizes with the genomically linked TN13 to membranes of the ER and the nuclear envelope

For subcellular localization studies, we transiently co-expressed TNT1-mYFP with a CFP-tagged ER marker (ER-ck; Nelson *et al*., 2007) in *Nbeds1a-1* mutant plants. Confocal laser scanning microscopy (CLSM) revealed that TNT1-mYFP, like TN13-mYFP, co-localizes with the ER-ck marker (Figure 3). The same localization was observed when expressing TNT1 or TN13 transiently under control of the respective *Np* (Figure S5). We analyze the subcellular localization of TN13-mYFP and TNT1-mYFP in *Arabidopsis* by using stable transgenic complementation lines expressing the respective mYFP fusion constructs under control of the *Np* or *35S* promoter (Figure S1). Fluorescence signals could not be detected for lines expressing the fusion protein constructs under control of the respective *Np*. For lines expressing TN13-mYFP or TNT1-mYFP under control of the *35S* promoter, we detected a fluorescence signal at the nuclear periphery and the ER (Figure S6). This is consistent with the observed localization in *N. benthamiana* and with the predicted N-terminal TM domains for TN13 and TNT1 (Figures 3, S3). Our data suggest that both TN13 and the genetically linked TNT1 localize to the ER membrane via their N-terminal TM domains.

**Figure 3.**
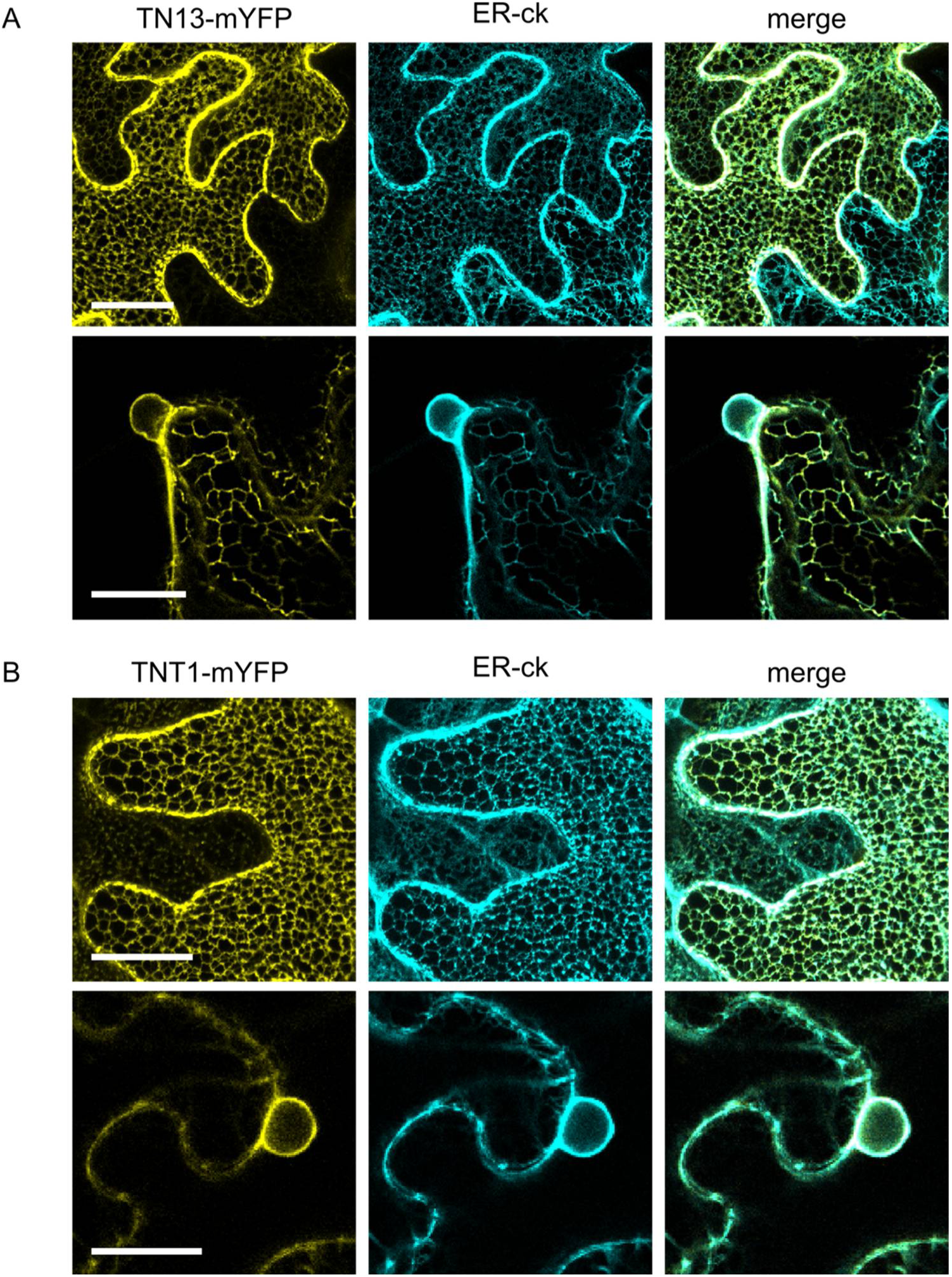
TN13-mYFP and TNT1-mYFP are ER membrane localized proteins. Localization of TN13 (A) and TNT1 (B) upon *Agrobacterium*-mediated transient expression in *Nbeds1a-1* leaves under control of the *35S* promoter. Confocal laser scanning microscopy was performed 2 days after *Agrobacterium* infiltrations. The top panels in (A) and (B) show z-stacks, the panels below show a single confocal plane crossing the nucleus. TN13-mYFP and TNT1-mYFP are shown in yellow, the co-expressed CFP-tagged ER marker (ER-ck, Nelson *et al*., 2005) is shown in cyan. Scale bar = 25 µm.

### TNT1-induced cell death depends on an atypical MHV motif

To investigate whether TNT1-triggered cell death in *N. benthamiana* depends on previously described functional NLR motifs, we conducted site-directed mutagenesis of the phosphate-binding loop (P-loop) and the MHD motif within the NB-ARC domain (Figure 4A). We found that the wild-type TNT1 protein contains an atypical MHV motif (Figures 4B). Mutations of MHD motifs to MHV have previously been shown to render NLRs auto-active (Bendahmane *et al*., 2002; Tameling *et al*., 2006; Williams *et al*., 2011). We surveyed a NLR dataset from 180 RefSeq reference plant proteomes (Toghani and Kamoun, 2024) and extracted MHD motifs of 56,686 NLRs to analyze the amino acid frequencies within the MHD motif. As expected, MHD constitutes the consensus sequence, and only a low number of NLRs (0.226 %) naturally contain a Valin (V) in the 3^rd^ position of the MHD motif (Figure 4B). In *Arabidopsis*, TNT1 is the only NLR containing an MHV motif (Supplementary Dataset 2). Mutating the wild-type MHV-motif of TNT1 to MHD or MHH abolished the cell death response upon transient expression in *N. benthamiana* wild-type plants, albeit protein accumulation was comparable to the TNT1 wild-type protein expressed in *Nbeds1a-1* (Figures 4, S7A). Mutagenesis of the P-loop in the wild-type TNT1 protein (containing the MHV motif) also abolished the cell death induction in wild-type *N. benthamiana* (Figures 4, S7B). We conclude that TNT1 requires the atypical MHV motif for the autoactive induction of cell death in a P-loop dependent manner.

**Figure 4.**
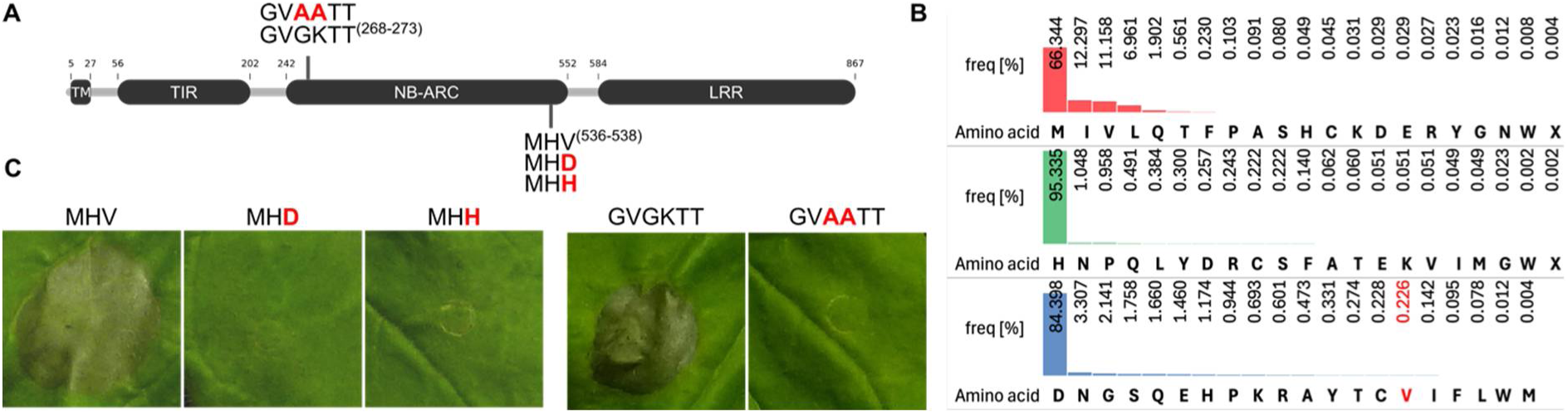
Cell death induction by TNT1 depends on the atypical MHV-motif and a functional P-loop. (A) Schematic representation of the TNT1 protein domain structure indicated by the amino acid positions within the TNT1 sequence. Amino acid exchanges introduced by site-directed mutagenesis are indicated in red. (B) Frequency of amino acids in each position of the MHD motif based on data of 56,686 NLRs from the RefSeq NLR database (Toghani and Kamoun, 2024; Ibrahim *et al*., 2024). X indicates ambiguous amino acid. (C) *Agrobacterium*-mediated transient expression of the indicated TNT1 wild-type and mutant variants fused to mYFP in *N. benthamiana* wild-type plants under control of the *35S* promoter. Cell death was photographed 5d after *Agrobacterium* infiltration.

### The TIR-domain of TNT1 is sufficient to trigger cell death in *N. benthamiana*

Expression of TIR domains was shown to be sufficient for cell death induction in *N. benthamiana* (Bernoux *et al*., 2011; Nandety *et al*., 2013; Williams *et al*., 2014; Nishimura *et al*., 2017). To analyze if the TNT1 TIR domain is sufficient for cell death induction, we expressed individual TNT1 domains, or truncated variants in wild-type *N. benthamiana* as mYFP-tagged fusion proteins.

Expressing TNT1 without the predicted TM domain (ΔTM, TIR-NB-LRR-mYFP) failed to induce cell death (Figures 5, S8A), and CLSM shows that this variant is no longer ER localized, but shows a cytoplasmic localization (Figure 5). Microsomal fractionation, followed by immunoblot analysis, further revealed that the ΔTM variant partially lost association with the endomembrane system, in contrast to the wild-type variant containing the predicted TM domain (Figure S8B). Since the LRR domain can have an auto-inhibitory effect on HR induction (Slootweg *et al*., 2013; Schreiber *et al*., 2016), we expressed a construct lacking both, the TM and LRR domain (TIR-NB-mYFP). Removal of the LRR domain from the ΔTM variant shifted the localization from exclusively cytoplasmic to nucleocytoplasmic, and induced a cell death response (Figures 5, S8A and B). Cell death was also induced upon expression of the ER-associated TM-TIR-NB-mYFP and TM-TIR-mYFP, or a TIR-mYFP variant that showed strong nuclear accumulation (Figures 5, S8A and B). While the TM-TIR-NB-mYFP and TM-TIR-mYFP variants are predominantly found in the endomembrane fraction, the TIR-mYFP fusion is predominantly soluble (Figure S8B). The NB-mYFP and LRR-mYFP proteins are nucleocytoplasmic or exclusively cytoplasmic, whereas the expression of an mYFP fusion to the predicted TM domain, followed by the C-terminal flanking region (amino acids 1-43), is sufficient to anchor the protein to the ER-membrane (Figure 5, S8A and B; Lee *et al*., 2011). No cell death was visible after expression of the NB, LRR- or TM-mYFP fusions (Figure 5).

**Figure 5.**
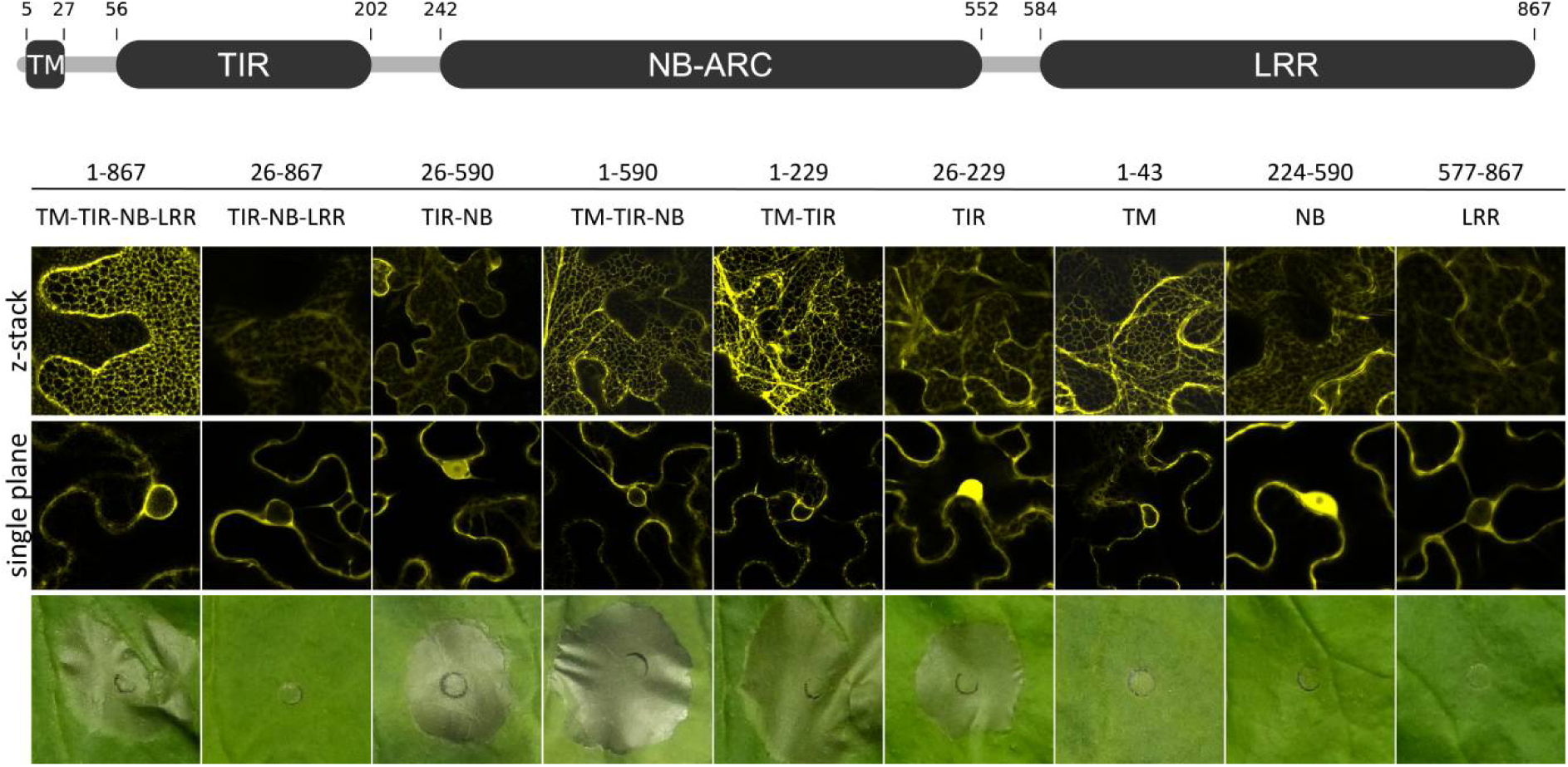
The TIR domain of TNT1 is sufficient for cell death induction. A schematic overview of the predicted TNT1 protein domain structure, with beginning and end of the respective domains indicated by amino acid positions at the top. The amino acids and domains of TNT1 expressed as mYFP fusions are indicated above CLSM images showing z-stacks (upper panel), and single confocal planes (middle panel) crossing the nucleus. Confocal images were taken 2 d after *Agrobacterium*-infiltration of *Nbeds1a-1* leaves. Transient expression for cell death assays (bottom panel) was performed in leaves of *N. benthamiana* wild-type plants, infiltrated leaf areas were photographed 5 d after *Agrobacterium* infiltration. Confocal images showing the localization of the full length TNT1 protein are also displayed in Figure 3B.

Since the expression of truncated TNT1 versions influenced the subcellular localization and ability to trigger cell death, we also analyzed domain truncations and single domains of the TN13 protein for their localization and ability to induce cell death. Expression of a ΔTM variant (TIR-NB-mYFP) completely abolished the localization of TN13 to the ER, but not its endomembrane association (Figure S8C-E). Whereas TM-TIR-mYFP or TM-mYFP variants showed an ER-localization, the TIR-mYFP as well as the NB-mYFP variants, did not localize to the ER, but remained partially associated with the endomembrane system (Figure S8C-E). None of the TN13 protein variants induced cell death upon transient expression in *N. benthamiana* wild-type plants (Figure S8C).

In summary, expression of the TNT1 TIR domain is sufficient to trigger a cell death response in *N. benthamiana*. The TM domain of TNT1 and TN13 is required and sufficient for the localization to the ER membrane. An ER localization of the full length TNT1 protein, but not of the TIR domain-containing truncated TNT1 protein variants, appears to be required for cell death induction (Figure 5), suggesting TM-specific suppressive effects of the LRR domain on cell death execution in *N. benthamiana*.

### The predicted catalytic glutamate residue of the TNT1 TIR domain is required for cell death induction in *N. benthamiana*

A conserved catalytic glutamate (E) residue within TIR domains is required for degradation of NAD^+^/NADP^+^, leading to accumulation of cyclic ADP-ribose variants (v-cADPR) and cell death in plants (Horsefield *et al*., 2019; Wan *et al*., 2019; Maruta *et al*., 2022). The TNT1 and the TN13 TIR domains both contain conserved glutamate residues at amino acid position 136 and 139, respectively (Figure, S3A). Consistent with its proposed function, mutagenesis of the TNT1 glutamate to alanine (E_136_A) abolished cell death induction upon transient expression of full length TNT1, TM-TIR or TIR-only variants in *N. benthamiana* wild-type plants (Figures 6, S8F). In line with our finding that TNT1 requires the EDS1-SAG101-NRG1 module, this result suggests that TNT1 induces cell death via TIR domain enzymatic activity.

**Figure 6.**
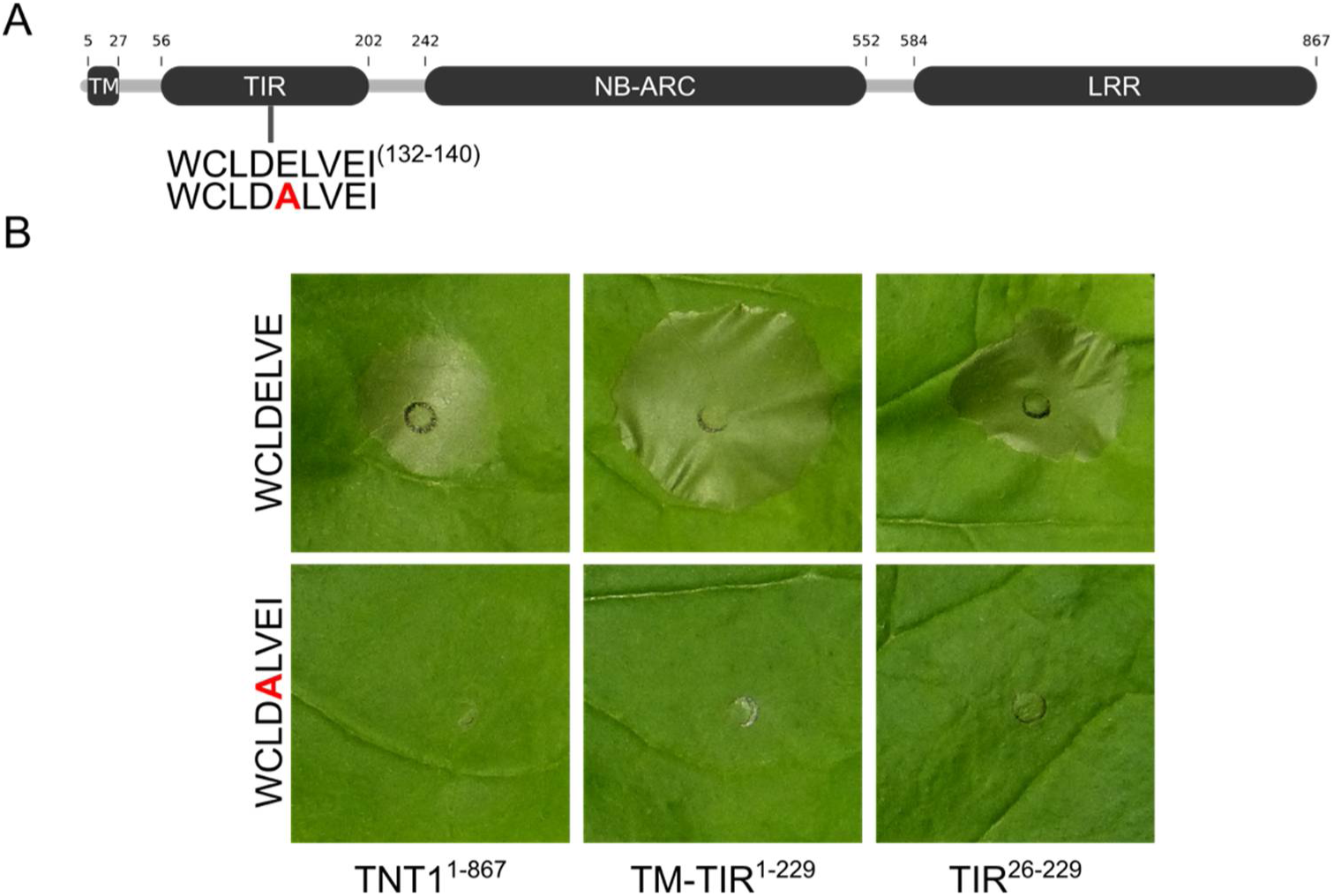
Cell death induction of TNT1 in *N. benthamiana* wild-type plants depends on the predicted catalytic glutamate residue of the TIR domain. (A) Schematic representation of the TNT1 protein domain structure. The predicted beginning and end of the respective protein domains are indicated by amino acid positions at the top. Amino acids 132-140 of the predicted catalytic motif in the TIR domain, containing the essential catalytic glutamate residue (E_136_), are depicted in single letter code. The amino acid exchange introduced via site-directed mutagenesis is indicated in red (A and B). (B) *Agrobacterium*-mediated transient expression of the indicated TNT1 wild-type and E_136_A mutant variants in *N. benthamiana* wild-type plants under control of the *35S* promoter. Cell death was photographed 5d after *Agrobacterium* infiltration.

### The genomic region encoding *TN13* and *TNT1* was duplicated and encodes NLRs that interact with TN13 and TNT1

The co-expression of TIR domains from TIR-type NLR pairs can suppress cell death induction of executor NLR TIR domains when expressed on their own (Williams *et al*., 2011; Huh *et al*., 2017). Since the overexpression of TNT1-mYFP in *Arabidopsis* does not induce autoimmunity, but triggers cell death in *N. benthamiana* (Figures 2, S2), we speculated that TNT1, TN13 and/or additional TNLs might interact with each other to suppress cell death induction in *Arabidopsis*. However, transient co-expression of TN13-mYFP and TNT1-mYFP in *N. benthamiana* did not results in a suppression of cell death (Figure S9).

*Arabidopsis* chromosome 3 does not encode for other NLRs within close chromosomal proximity to the *TN13*/*TNT1* pair (Meyers *et al*., 2003), however, the genomic region encoding *TN13* and *TNT1* shows persistence of *NLR* gene presence in a segmentally duplicated chromosomal region (Simillion *et al*., 2002; Meyers *et al*., 2003). This chromosomal region between *AT5G14060* and *AT5G18490* on chromosome 5 was duplicated segmentally from *AT3G01015* to *AT3G04350* on chromosome 3 (Figure S10, Meyers *et al*., 2003). The duplicated region of chromosome 5 contains an expanded number of TIR-type NLR genes, consisting of an uncharacterized genetic singleton (*AT5G17680*) and two NLR clusters (Figure 7A; based on the cluster definition by Meyers *et al*. (2003)). The first cluster encodes the head-to-head sensor-executor pair *CHS3*/*CONSTITUTIVE SHADE-AVOIDANCE1* (*CSA1*) (*AT5G17890/80*) which guard BAK1 and BIR3 (Schulze *et al*., 2022) and an uncharacterized TNL (*AT5G17970*). The second cluster contains *DOMINANT SUPRESSOR OF CAMTA3 NUMBER2* (*DSC2*, *AT5G18370*) that is required for the *camta3* autoimmune phenotype (Lolle *et al*., 2017), *HOPB-ACTIVATED RESISTANCE1* (*BAR1*, *AT5G18360*) that recognizes the *Pseudomonas* effector HopB1 (Laflamme *et al*., 2020), and an uncharacterized *TNL* (*AT5G18350*) in tandem orientation (Figure 7A). Phylogenetically, AT5G18350 and DSC2 are the closest relatives of TNT1 and TN13 within the duplicated chromosomal region (Figures 7B). In addition to the close phylogenetic relationship of DSC2 to TNT1, DSC2 also contains a predicted N-terminal transmembrane domain and localizes to the ER when transiently expressed in *Nbeds1a-1* plants (Figures 7B, S11, S12). The other TIR-type NLRs encoded in the duplicated region show a nuclear, cytoplasmic or nucleocytoplasmic distribution in *Nbeds1a-1* (Figure S12). Expression of all NLRs from the segmentally duplicated region on chromosome 5 triggered a cell death response in wild-type *N. benthamiana* plants, except for CHS3 (Figure S13). Co-expression of the NLRs together with TNT1 did not suppress cell death induction (Figure S14).

**Figure 7.**
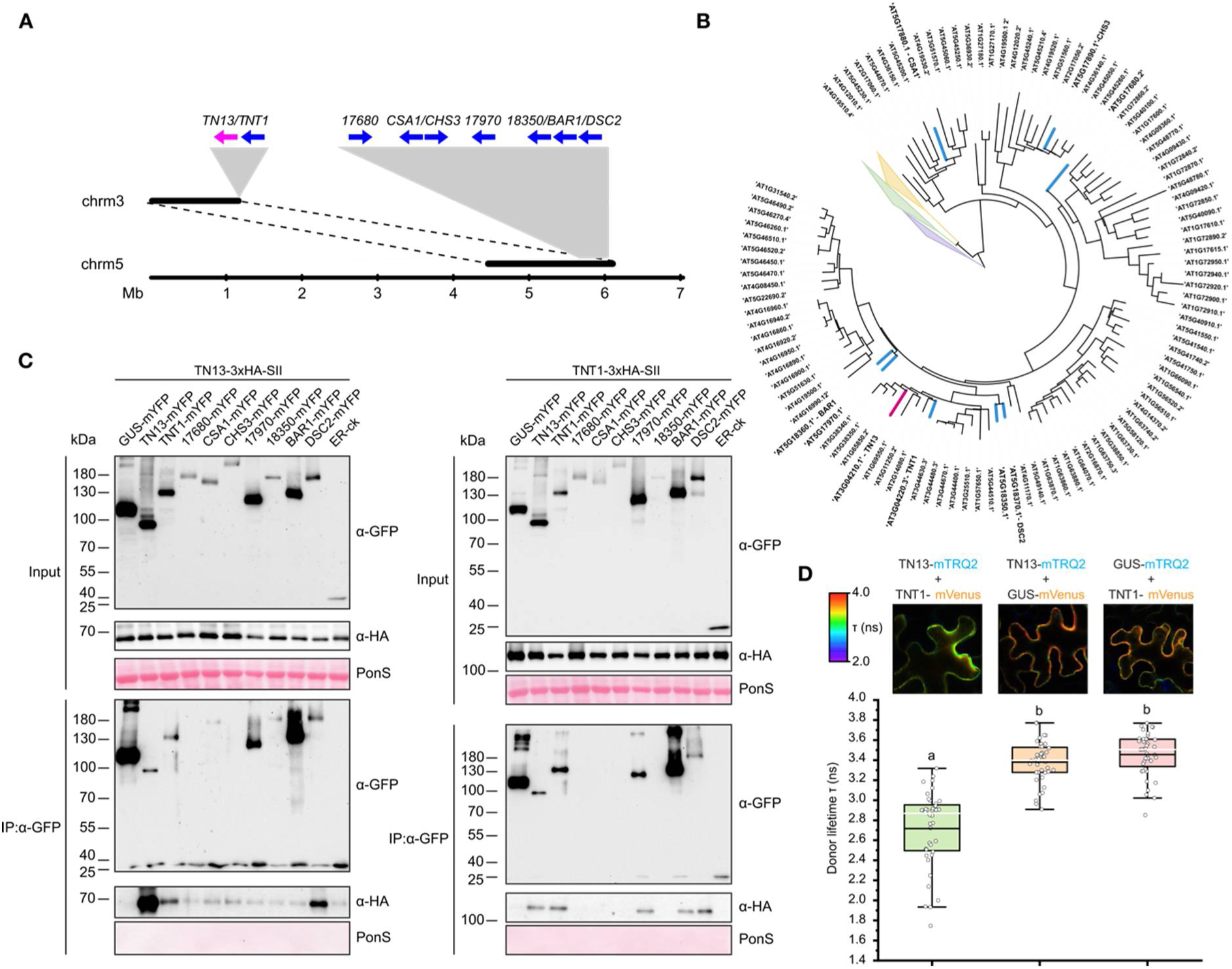
TN13 and TNT1 interact with each other and with NLRs encoded by a segmentally duplicated genomic region on *Arabidopsis* chromosome 5. (A) Schematic representation of the position of the segmental duplication of chromosome (chrm) 3 and chromosome 5. Full length *TIR*-type *NLRs* are depicted as blue arrows, *TN*-type *NLRs* in magenta. The corresponding gene names or loci are indicated. (B) Phylogenetic tree of *Arabidopsis* NLRs based on the NB-ARC domain. RNL (purple), CCG10 (yellow), and CNL (green) clades are collapsed. (C) mYFP-tagged proteins were immunoprecipitated from protein extracts obtained 2 days after *Agrobacterium*-infiltration for transient expression of the indicated fusion proteins in *Nbeds1a-1* plants, using GFP-Trap magnetic agarose beads (IP: α-GFP). Co-immunoprecipitation of 3xHA-StrepII (3xHA-SII)-tagged NLRs was detected by α-HA immunoblots. Total protein extracts (Input) were probed with α-GFP and α-HA antibodies, respectively. GUS-mYFP and a CFP-tagged ER marker (ER-ck; Nelson *et al*., 2007) served as negative controls. All samples were separated on 7.5 % SDS polyacryamide gels, blotted onto nitrocellulose membranes and probed with the indicated primary antibodies. Horseradish peroxidase coupled secondary antibodies were used for detection. PonceauS (PonS) staining of the membranes was used as loading control. (D) FLIM measurements of transiently expressed donor proteins TN13-mTRQ2 or GUS-mTRQ2 in *N. benthamiana eds1a-1* epidermal cells, two days post-infiltration (dpi) of *Agrobacteria* for co-expression of the fusion proteins indicated on top. Donor lifetime heat maps of representative cell areas used for FLIM measurements are shown according to the scale on the left. Data are presented as box plots. Dots represent individual measurements, the lower and upper limits of the box indicate the interquartile range, and the black and white lines represent the mean and median, respectively. Different letters above the plots indicate a statistically significant difference among different groups (n=40, one-way ANOVA, Tukey’s test, P<0.001).

To investigate potential NLR complex formations, we conducted co-immunoprecipitation experiments after co-expression of TN13-mYFP, or TNT1-mYFP, together with 3xHA-SII tagged candidate NLRs of the duplicated region in *Nbeds1a-1* plants (including 3xHA-SII tagged versions of TN13 and TNT1 to monitor potential TN13 and TNT1 homomeric associations). Co-detection of 3xHA-SII tagged NLRs, after immunoprecipitation of mYFP-tagged proteins shows that TNT1, but not TN13, co-immunoprecipitates BAR1 and the uncharacterized NLR encoded by *AT5G17970* (Figure 7). In addition, both TN13 and TNT1 can form homo-oligomers (Figure 7C) as well as hetero-oligomers with each other, and both proteins interact with DSC2 (Figure 7C). GUS-mYFP and the CFP-tagged ER marker (ER-ck) were used as a negative control (Figure 7). As the immunoprecipitation via the C-terminal mYFP tag was inefficient for CHS3 and CSA1, as well as for the uncharacterized NLRs encoded by *AT5G17680* and *AT5G18350*, no conclusion on their interaction with TN13 and TNT1 can be made. To independently investigate the results obtained by Co-IP for the interaction between TN13 and TNT1, we employed a CLSM-based Förster resonance energy transfer-fluorescence lifetime imaging (FLIM-FRET) analysis using TN13-mTRQ2 as donor and TNT1-mVenus as acceptor for fluorescence transfer. Co-expression of TN13 with TNT1 significantly reduced the fluorescence lifetime of TN13-mTRQ2, indicating *in planta* interaction (Figure 7D). A reduction in lifetime was not observed when expressing TN13-mTRQ2 together with GUS-mVenus, used as a negative control (Figure 7D). The identified interactions and our subcellular localization studies suggest that TN13 and TNT1, together with NLRs from the segmentally duplicated cluster on chromosome 5, might form a larger TIR-type NLR immune receptor network at the cytoplasmic side of the ER-membrane and the nuclear envelope in *Arabidopsis* cells.

## DISCUSSION

NLRs can engage in intermolecular interactions within pairs or networks (Le Roux *et al*., 2016; Sarris *et al*., 2015; Césari *et al*., 2014; Wu *et al*., 2017; Contreras *et al*., 2023). An additional level of complexity in TIR-type NLR function is the requirement of EDS1 family proteins and RNL-type helper NLRs for downstream signaling (Castel *et al*., 2019; Wu *et al*., 2019; Saile *et al*., 2020; Wu *et al*., 2024; Yu *et al*., 2024). Recent examples also show the functional requirement of truncated TIR-type NLRs in immunity. Truncated TIR-NLRs were proposed to act as sensors that interact with and require an executor TIR-type NLR for efficient downstream signaling (Zhao *et al*., 2015; Zhang *et al*., 2017; Tong *et al*., 2017; Liang *et al*., 2019), albeit direct contributions to immune signaling have also been demonstrated (Staal *et al*., 2008; Nandety *et al*., 2013; Nishimura *et al*., 2017).

### TN13 and TNT1 are required for full immunity to mildly virulent *P. syringae*

Our analysis revealed that *TNT1*, like *TN13*, is required for full resistance to *Pst* DC3000 (ΔAvrPto/AvrPtoB), and co-localizes with TN13 to the ER and perinuclear membrane (Figures 1, 3, S6; Roth *et al*., 2017). TN13 and TNT1 form homo-oligomers and interact with each other when co-expressed in *N. benthamiana* (Figures 7C). This suggests that TN13 and TNT1 may form an NLR pair involved in regulating defense responses at the ER. Since both *NLRs* are genetically required for full resistance to mildly virulent *Pst* (Figure 1), TN13 and TNT1 may recognize *Pst* effector(s), and initiate immune signaling to restrict growth of virulent bacteria (Thomma *et al*., 2011). Yeast two-hybrid assays by Nandety *et al*. (2013) suggest that TN13 interacts with the *Pst* effector HopY1 and the nematode effector Rbp001. This would be consistent with a potential sensor function of TN13. Notably, BAR1 is required for recognition of the *Pst* effector HopB1 (Laflamme *et al*., 2020). Since BAR1 interacts with TNT1 in *N. benthamiana* (Figure 7C), TNT1 might also be involved in regulating HopB1-triggered defense responses.

The transient expression of TNT1 in wild-type *N. benthamiana* triggers a cell death response that depends on *EDS1/SAG101/PAD4*, whereas no cell death is induced by TN13 (Figures 2, S8C, S9). We hypothesize that TNT1 could function as an executor NLR for TN13, which could act as a sensor lacking the capacity to produce small molceules for immune signaling on its own. Examples of TN-type sensor NLRs include CHS1 and TN2, which both monitor the cellular homeostasis of the membrane-associated E3 ubiquitin ligase SAUL1, and signal independently via the executor TNL SOC3 (Zhang *et al*., 2017; Tong *et al*., 2017; Liang *et al*., 2019).

### The transmembrane domains of TN13 and TNT1 are required for ER localization

TN13 and TNT1 both contain N-terminal transmembrane (TM) domains that are required and sufficient to localize the mYFP-tagged fusion proteins to the ER membrane (Figures 5, S3, S6, S8A-E). Multiple alleles of the TNL RPP1 from different *Arabidopsis* accessions also contain a hydrophobic extension N-terminal of the TIR domain (Schreiber *et al*., 2016). RPP1-WsA localizes to ER and/or Golgi membranes, and the N-terminal extension is partially required for membrane localization and function in immunity (Weaver *et al*., 2006). Deleting the TM-domain of TNT1 lead to a loss of cell death when expressing this protein in *N. benthamiana* (Figure 5), which was re-established by additional deletion of the LRR domain. This probably reflects an inhibitory function of the LRR on TIR domain activity, as previously described for other NLRs (Figure 5; Slootweg *et al*., 2013; Ntoukakis *et al*., 2013; Schreiber *et al*., 2016). As an ER membrane localization of TIR-containing TNT1 truncations is not essential for cell death induction, the ER membrane association might represent an additional mechanism for regulating the activation of the full length NLR that otherwise would be suppressed by the LRR domain (Figure 5). TNT1 cell death induction in *N. benthamiana* depends on SAG101b/NRG1, but not on PAD4*/*ADR1 (Figure 2). Albeit such functional cooperativity is consistent with the finding that *Arabidopsis* NRG1A and NRG1B also localize to the ER (Wu *et al*., 2019; Ibrahim *et al*., 2024), the detailed functional interplay between TNL activation and NRG1 resistosome formation at the ER remains to be explored.

### Cell death induction by TNT1 requires an atypical MHV motif

The predicted catalytic glutamate residue within the TIR domain of TNT1 is required for cell death induction in *N. benthamiana* (Figure 6), suggesting that TNT1 might degrades NAD^+^/NADP^+^ to produce small molcules for the induction of immunity, as previously described for TIR domains (Wan *et al*., 2019; Horsefield *et al*., 2019). The TIR domains of the paired sensor TNLs CHS3 and RRS1 lack the catalytic glutamate residue, enzymatic activity, and do not induce cell death in *N. benthamiana* when expressed on their own (Figure S16; Wan *et al*., 2019; Horsefield *et al*., 2019), whereas the TN-type sensor CHS1 contains the catalytic glutamate residue but does not induce cell death upon expression in *N. benthamiana* (Liang *et al*., 2019). Likewise, TN13 also contains the predicted catalytic glutamate (Figures S3A, S9, S11), but expression of TN13 in *N. benthamiana* does not trigger a cell death response (Figures 2A, S8C). TN2 induced cell death in *N. benthamiana* is suppressed by co-expression of EXO70B1 (Wang, Liu, *et al*., 2019). The RRS1-R/RPS4 sensor/executor pair, forms a higher order oligimeric complex in their inactive state, which likely requires reconfiguration upon effector interaction to derepress the RPS4 TIR domain and enable NADase ezyme function (Ahn *et al*., 2025). It therefore remains possible that cell death induction upon expression of TN13 in *N. benthamiana* requires an additional executor as NLR signaling partner, or that cell death induction is suppressed via association with an interaction partner present in *N. benthamiana*. This might also explain why a stunted growth phenotype can be observed in the TN13 *Arabidopsis* overexpression lines, albeit no cell death is visible when expressing TN13 in *N. benthamiana* (Figure S2).

TIR domain catalytic activity depends on homo-oligomerization to form a holoenzyme resistosome (Horsefield *et al*., 2019). Both TN13 and TNT1 show homo and hetero-oligomer formation (Figure 7). Remarkably, the TNT1 protein contains an MHV motif within the NB-ARC domain that diverges from the canonical MHD motif in other TNLs and is required for cell death induction (Figures 4). TNT1 is the only *Arabidopsis* TNL containing an MHV motif within the NB-ARC domain of the wild-type protein. The presence of the MHV motif may confer higher affinity for ATP, rather than ADP, and constitutively shift TNT1 to an active state (Sukarta *et al*., 2016). TNT1 would thus require efficient intra-or/and intermolecular repression to avoid inadvertent cell death signaling in the absence of a pathogen trigger. The CHS3/CSA1 sensor/executor pair was shown to form heterodimers and oligomerization and cell death singaling is suppressed by the receptor kinases BAK1 and BIR3, which are guarded by CHS3/CSA1 (Yang *et al*., 2022; Schulze *et al*., 2022; Yang *et al*., 2024). In a similar scenario, TNT1 could function as an executor that interacts with sensors, such as TN13, and could potentially be suppressed by an unknown guardee in *Arabidopsis*, but not upon transient expression in *N. benthamiana*.

The MHV motif containing TNT1 could therefore function as an integrative hub for HR induction at the interface of the ER and the cytoplasm, the PM or other organelles, that is activated upon derepression. It has previously been reported that expression of *TNT1* is upregulated following elicitor treatment, *Pseudomonas* infection, or in autoactive mutant plants exhibiting constitutive defence responses (Tian *et al*., 2021; Lang *et al*., 2022). In addition, upregulation of *TNT1* expression is dependent on MPK3/6 kinase signaling and induced expression of *TNT1* activates salicylic acid biosynthesis genes (Lang *et al*., 2022). Therefore, TNT1 might also contribute to the induction of basal defense which would explain the defects of the *tnt1* mutant in basal resistance to *Pst* DC3000 (ΔAvrPto/AvrPtoB).

### TN13 and TNT1 interact with each other and NLRs in a segmentally duplicated genomic region

The region of chromosome 3 containing *TN13*-*TNT1* was duplicated segmentally to chromosome 5 (Figures 7, S10; Simillion *et al*., 2002; Meyers *et al*., 2003). The duplicated region on chromosome 5 contains syntenic regions with an expanded number of *NLR* genes (Meyers *et al*., 2003, Figures 7, S10). Phylogenetic analysis of the complete repertoire of *Arabidopsis* (Col-0) TIR-type NLRs suggest that TN13 and TNT1 are closely related to NLRs of the tandem cluster, including AT5G17970 (Figure 7; Van de Weyer *et al*., 2019). This is expected based on a model of conserved synteny, where physical proximity correlates with close phylogenetic relationship (Baumgarten *et al*., 2003) and would further support that the tandem cluster components *AT5G18350*/*BAR1*/*DSC2*, and potentially *AT5G17970*, originated from *TN13* and/or *TNT1*. It has been proposed that tandem gene duplications play a major role in the evolution of plant NLRs, while segmental duplications are less prevalent, but partially explain the expansion of *NLRs* within the *Arabidopsis* genome to distinct chromosomal regions (Richly *et al*., 2002; Leister, 2004; Cannon *et al*., 2004).

The duplication of *NLR*s can be considered as a mechanism to generate diversification, to produce new NLR variants for effector detection (Leister, 2004; Cannon *et al*., 2004; Barragan and Weigel, 2021). All *NLRs* within the segmentally duplicated clusters of chromosome 3 and 5 belong to the *Arabidopsis* core NLRome, with the exceptions of *TN13*, *AT5G17970* and *AT5G18350* that are part of the shell NLRome and potentially arose via tandem duplication (Figure S14; Van de Weyer *et al*., 2019). Expanded tandem duplicated regions, such as the *RPP1* cluster, were also identified as *DANGEROUS MIX* (*DM*) loci. These loci can lead to hybrid necrosis upon crossing of different accessions due to NLR repertoire incompatibility caused by diversifying pressure on loci in the *NLR* clusters (Bomblies *et al*., 2007; Chae *et al*., 2014; Alcázar *et al*., 2014; Stuttmann *et al*., 2016; Atanasov *et al*., 2018).

Both TN13 and TNT1 interact with DSC2 (AT5G18370). In addition, TNT1 interacts with BAR1 (AT5G18360) and the uncharacterized TNL protein encoded by *AT5G17970* (Figure 7). These findings suggests that TN13 and TNT1 could be part of a larger TNL signaling network that involves TNLs from the duplicated cluster. The hetero-pairing of TN13 and TNT1 with NLRs of the duplicated cluster on chromosome 5 might be explained by their close phylogentic relation. In fact, several sensor-executor NLR pairs that are genomically linked fall into neighboring phylogenetic sub-clades (Césari *et al*., 2014; Van de Weyer *et al*., 2019; Ahn *et al*., 2025). Since *tn13* mutation as well as *TNT1* over-expression in stable transgenic plants did not show an auto-immune phenotype (Figure S2), we speculate that TNT1 is repressed by additional TNLs or other host proteins. Most NLRs encoded in the duplicated cluster contain a predicted catalytic glutamate residue within the TIR domain and trigger a cell death response when expressed individually, or together with TNT1, in *N. benthamiana* (Figures S12, S13). Whether TNLs of the duplicated region function together with TNT1 or potentially influence cell death induction of TNT1 upon co-expression of multiple TNL combinations, remains to be investigated.

## EXPERIMENTAL PROCEDURES

### Plant materials

All plants were grown in environmentally controlled growth chambers with 65 % relative humidity. *Arabidopsis thaliana* plants were grown on soil for infection studies, or on ½ MS agar-plates for microscopy analyses, under an 8/16 h light/dark regime at 22/18°C. *Nicotiana benthamiana* plants were grown under a 16/8 h light/dark regime at 25/22°C. The *N. benthamiana eds1a-1* mutants (Ordon *et al*., 2017), *pad4* and *sag101a*/*b* (single, double and triple) mutant combinations (Gantner *et al*., 2019; Lapin *et al*., 2019), as well as the *Arabidopsis tn13* (Roth *et al*., 2017)*, snc1* (Li *et al*., 2001) and Col *eds1*-2 (Bartsch *et al*., 2006) mutants were previously described. The *N. benthamiana adr1 nrg1* and *nrg1* mutant lines were kindly provided by Jane Parker (MPIPZ, Cologne) and Johannes Stuttmann (BIAM - CEA Cadarache). The *tnt1-1* mutant line were obtained from the Nottingham Arabidopsis Stock Centre (NASC, http://arabidopsis.info). Homozygous mutants were isolated via PCR-based genotyping, using primers listed in Table S1. Stable transgenic *Arabidopsis* plants were generated by using the floral dip method (Clough and Bent, 1998).

### Plasmid constructions

The full-length genomic sequences (with or without promoter region) were PCR amplified without stop codon, using gene specific primers listed in Table S1, and cloned into pENTR/D-TOPO (ThermoFisher Scientific, https://www.thermofisher.com) Gateway entry vectors. The generation of truncations, or site-directed mutagenesis, was conducted by sub-cloning into pENTR/D-TOPO and/or mutagenesis PCR (*Q5^®^* Site-Directed *Mutagenesis* Kit, NEB), using primers listed in Table S1 and respective pENTR/D-TOPO entry clones as templates. The pENTR/D-TOPO *GUS*+*intron* and *TN13* entry vectors (Roth *et al*., 2017), the *ER-ck* (Nelson *et al*., 2007), and *NRG1* (Lapin *et al*., 2019) expression vectors were described previously. All expression constructs were generated via Gateway LR-reactions with pXCSG-GW-mYFP, pXCG-GW-mYFP or pXCSG-GW-3xHA-StrepII destination vectors (Witte *et al*., 2004; Feys *et al*., 2005) and respective pENTR/D-TOPO sequence-verified entry vectors. All generated expression vectors were transformed into *Agrobacterium tumefaciens* strain GV3101 pMP90RK (Koncz and Schell, 1986), using electroporation.

### *Pseudomonas* infection assays

A bacterial suspension of *Pst* DC3000 (ΔAvrPto/AvrPtoB; Lin and Martin, 2005) with a density of 1 × 10^5^ cfu ml^−1^ in 10 mM MgCl_2_ was infiltrated into rosette leaves of five 5-weeks-old soil grown *Arabidopsis* plants, using a needleless syringe. Colony forming units were determined 1 h (d0) and 3 days (d3) after infiltration.

### Transient expression in *N. benthamiana*

*Agrobacterium* overnight cultures were harvested by centrifugation, resuspended in infiltration buffer (10 mM MgCl_2_, 10 mM MES pH 5.5, 150 µM acetosyringone), and incubated for 2-3 h at room temperature. For co-expressions, the bacterial suspension was adjusted to an OD_600_ of 0.3 for each strain and infiltrated into the abaxial side of 4-5 weeks old *N. benthamiana* leaves, using a needleless syringe. An *Agrobacterium* strain expressing the silencing suppressor p19 was co-infiltrated in all transient expression experiments. For HR cell death assays, the infiltrated leaf areas were photographed 5 days after infiltration. Confocal laser scanning microscopy was performed 2 days after *Agrobacterium* infiltration.

### Confocal laser scanning microscopy

For confocal microscopy, leaf discs of *Agrobacterium*-infiltrated *N. benthamiana* tissues or stable transgenic *Arabidopsis* plants were embedded in water. A Leica TSC-SP5 confocal laser-scanning microscope controlled by the Leica LAS AF software was used with a 20x/0.70 objective (PL APO, CS). An argon laser line at 514 nm was used for excitation of mYFP, and 458 nm for CFP. Emitted fluorescence was detected at 525 – 570 nm for mYFP, and 465 – 485 nm for CFP, using Leica HyD detectors. Chlorophyll auto-fluorescence was detected at 680–720 nm, using a Leica PMT detector. All channels were scanned sequentially. ImageJ was used to merge channels (Schindelin *et al*., 2012). FLIM-FRET experiments were conducted with transiently transformed *N. benthamiana* leaves using a Leica TCS SP8 microscope equipped with the FALCON time-correlated single photon counting system. The mTurquoise2 (mTRQ2)-tagged donor proteins were excited using a pulsed diode laser at 440 nm, and the mVenus-tagged acceptor proteins were excited with a pulsed white light laser at 514 nm. A pulse rate of 40 kHz and a two-channel sequential excitation were applied. The fluorescence emission was detected at 465-505 nm for mTRQ2 and 520-560 nm for mVenus using HyD SMD detectors. Images were acquired when at least 100 photons/pixel were collected in the brightest channel. The picture format was set to 512 x 512 pixels. Selection of regions of interest and FLIM data fitting were performed using the LAS X Single Molecule Detection software module (v3.5.5). For fluorescence lifetime calculations, a mono-exponential re-convolution fitting model was applied. Statistical analysis and plotting of the donor lifetime data was performed in Origin 2020 (OriginLab Corp., Northampton, USA).

### Total protein extracts, co-immunoprecipitation, microsomal fractionation and immunoblot analysis

To generate total protein extracts from *Arabidopsis* samples, leaf tissues were frozen in liquid nitrogen, homogenized and mixed with protein extraction buffer (250 mM sucrose, 100 mM HEPES-KOH (pH 7.5), 5 % (v/v) glycerol, 2 mM Na_2_MoO_4_, 25 mM NaF, 10 mM EDTA, 1 mM DTT, 0.5 % (v/v) Triton X-100, plant protease inhibitor cocktail (#P9599, Sigma)), cell debris was removed by centrifugation and filtration through a 95 µm nylon mesh. Protein concentrations were quantified using Bradford reagent (Biorad). Protein samples were adjusted to equal concentrations, mixed with 4x SDS-PAGE loading dye (250 mM Tris-HCl (pH 6.8), 8 % (w/v) SDS, 40 % (v/v) Glycerol, 0.04 % (w/v) Bromophenol blue, 400 mM DTT) and boiled. For co-immunoprecipitation analysis (Co-IP) after *Agrobacterium*-mediated transient expression in *N. benthamiana*, 7.5 µl GFP-trap magnetic agarose beads (Chromotek) were equilibrated in protein extraction buffer, and added to each total protein extract generated as described for *Arabidopsis* samples, without adjusting total protein concentrations. Incubation was performed for 3 h at 4 °C under constant rotation before the magnetic agarose beads were isolated and washed 3 times in 1 ml extraction buffer, using a magnetic rack. Proteins were eluded by boiling in 4x SDS loading dye. For microsomal fractionation after *Agrobacterium*-mediated transient expression in *N. benthamiana*, total protein extracts were prepared as described for *Arabidopsis* samples, without adjusting total protein concentrations, using a buffer without Triton X-100. Total protein extracts were subjected to ultra-centrifugation at 100.000 *g*, and an aliquot of the supernatant (soluble fraction) was mixed with 4x SDS loading dye. The endomembrane pellet was washed 3 times in 1 ml extraction buffer and dissolved in extraction buffer containing 0.5 % (v/v) Triton X-100, before mixing with 4x SDS loading dye and boiling. For total protein extraction from *N. benthamiana*, leaf tissues were frozen in liquid nitrogen, homogenized and boiled in 2x SDS-PAGE loading dye. Cell debris was removed by centrifugation at 17.000 *g*, and the supernatant was used for SDS-PAGE and immunoblot analysis. All protein samples were separated on SDS polyacrylamide gels and transferred onto nitrocellulose membranes (Amersham Protran, 0.45 μm; GE Healthcare Life Sciences). Primary α-GFP (#11814460001, Roche, 1:5000), α-HA (H9658; Sigma-Aldrich, 1:5000), and α-PEPC (100-4163, Rockland, 1:7500) antibodies and secondary goat anti-mouse IgG-poly-HRP (polyclonal, #32230; ThermoFisher Scientific, 1:5000) or goat anti-rabbit IgG-poly-HRP (polyclonal, #32260; ThermoFisher Scientific, 1:5000) antibodies were incubated and detected using SuperSignal West Femto chemiluminescence substrate (#34095; ThermoFisher Scientific) on a ChemiDoc imaging system (BioRad).

### RNA isolation and RT-PCR analysis

Isolation of total RNA and RT-PCR analyses were conducted as described by Genenncher *et al*., (2016), using primers listed in Table S1 in a 20 µl reaction volume. HDGreen (Intas) was used for staining of the gel.

### DNA isolation and genotyping PCR

Isolation of total genomic DNA was performed using the CTAB method. Briefly, 100 mg of *Arabidopsis* leaf tissue was ground to a fine powder in liquid nitrogen using a TissueLyser (QIAGEN, Hilden, Germany). CTAB extraction buffer was added, mixed and incubatated for 20min at 65 °C. Chloroform was added and the aqueous phase was mixed with double volume Ethanol. The DNA was precipitated at −20 °C for 30 min and pelleted by centrifugation. DNA was disolved in ddH_2_O and standard PCR genotyping was performed using primers listed in Table S1. HDGreen (Intas) was used for staining of the agarose gel.

### Phylogenetic analyses

For phylogenetic analyses, the Araport11 protein sequences of the Col-0 NLRome (based on Lee and Chae, 2020) were retrieved by providing a list of the AGI locus IDs in the TAIR gene search function (https://www.arabidopsis.org/search/genes). Domain predictions were performed in Geneious Prime (Kearse *et al*., 2012) using InterProScan5 (Jones *et al*., 2014), Transmembrane domains were predicted using TMHMM (Krogh *et al*., 2001). The NB-ARC and Winged-Helix domains of NLRs were extracted from an aligment, based on annotations. The extracted domains were re-aligned and a phylogenetic tree was generated. Aligments were performed using MAFFT (Katoh *et al*., 2002; Katoh and Standley, 2013), FastTree (Price *et al*., 2010) was used to generate phylogenetic trees. An alignement of the NB-ARC domains and the phylogenetic tree file are available as Supplementary Dataset S3 under https://zenodo.org/records/15281087.

### Motif frequency analysis

For MHD motif frequency analysis an aligment of deduplicated NLRs from 180 RefSeq reference plant proteomes, containing only the NB-ARC domains (Toghani and Kamoun, 2024; Ibrahim *et al*., 2024), was manually trimmed to the MHD motif using Geneious Prime (Kearse *et al*., 2012). The trimmed file was used to detmerine the frequency for each amino acid in each position. The trimmed MHD motifs are available as Supplementary Dataset S2 under https://zenodo.org/records/15281087.

### Statistical analysis

All statistical analyses were performed using R, v 2024.12.1+563 or Origin 2020 (OriginLab Corp., Northampton, USA). A Kurskal-Wallis test, followed by Pairwise Wilcoxon tests with Benjamini-Hochberg correction were preformed on non-parametric data for pathogen infection assays after log transformation. For FRET-FLIM analysis, one-way ANOVA coupled with Tukey‘s test (p<0.001) was used on donor lifetime.

### Structural prediction

AlphaFold3 (Abramson *et al*., 2024) was used with a seed value of 1 to generate structural models of the TN13 and TNT1 TM-TIR domains. The predicted structures were analyzed in ChimeraX (Pettersen *et al*., 2021), where domains were colored according to their pLDDT scores. The PDB files of the structural models are available as Supplementary Dataset S1 under https://zenodo.org/records/15281087.

## ACKNOWLEGDEMENTS

We thank Johannes Stuttmann (BIAM - CEA Cadarache) and Jane Parker (MPIPZ, Cologne) for *N. benthamiana* mutant seeds. We also thank Jane Parker for the pXC(S)G Gateway destination vectors. We acknowledge Volker Lipka and Elena Petutschnig (Plant Cell Biology and Central Microscopy Facility of the Faculty of Biology and Psychology, University of Goettingen) for support and access to a confocal microscopy platform (Deutsche Forschungsgemeinschaft (DFG) grants INST 186/824-1 and INST 186/1277-1 to V.L.). We are also grateful to Xin Li (UBC, Vancouver) for useful discussion. We thank AmirAli Toghani (TSL, Norwich) for providing an alignment of NLR NB-ARC domains of a RefSeq NLR database. This research was funded by the DFG IRTG 2172 “PRoTECT” program of the Goettingen Graduate Center of Neurosciences, Biophysics, and Molecular Biosciences and the DFG research grant WI 3208/4-2 to M.W.

## SUPPORTING INFORMATION

**Figure S1.**
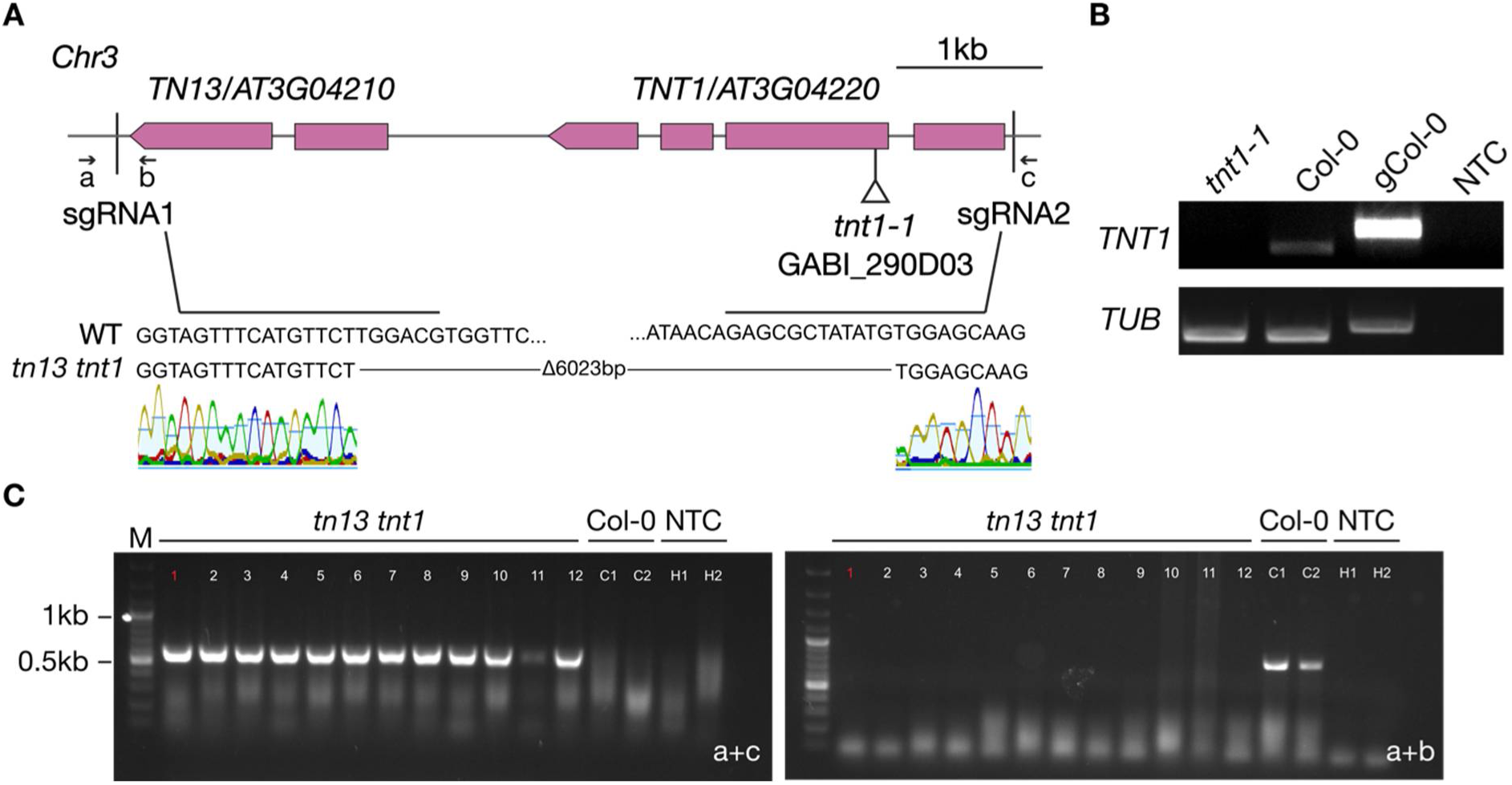
Genotyping of *tnt1-1* and *tn13 tnt1* mutant lines. (A) Gene structures of genetically linked *TN13* and *TNT1* drawn to scale. Exons are represented as boxes and introns as solid lines. The positions of the T-DNA insertion in *tnt1-1* is marked by a triangle below the gene structure. The position of single guide RNAs (sgRNA) used to generate the *tn13 tnt1* double mutant are indicated. Flanking (a+c) and gene specific (a+b) primers used for genotyping of the *tn13 tnt1* double mutant are indicated. (B) RT-PCR analysis with cDNAs, transcribed from RNA of four weeks old *tnt1-1* T-DNA insertion lines, grown on soil under short day conditions. *TUBULIN* (*TUB*) was used as control. PCR products were separated by agarose gel electrophoresis and stained by HDGreen. (C) PCR-genotyping of the *tn13 tnt1* double mutant. Standard PCR was performed with primers flanking the single guide RNA (sgRNA) targets sites and gene specific primers to detect gene deletions. PCR products were separated by agarose gel electrophoresis and stained by HDGreen. C1/2 and H1/2 represent Col-0 and ddH_2_O controls, respectively. The red number above the band indicates the line used in subsequent experiments.

**Figure S2.**
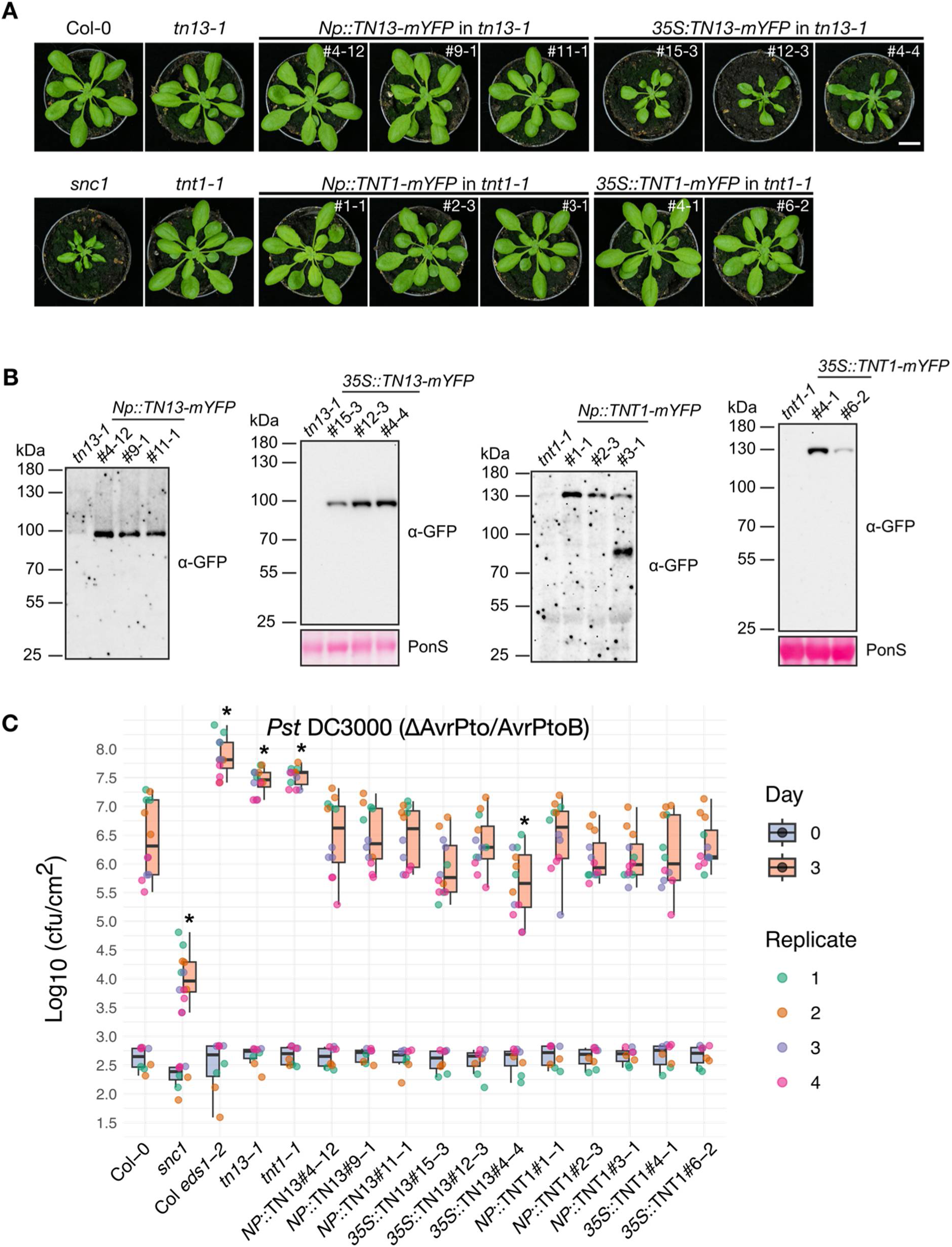
Phenotyping of *tn13* and *tnt1* mutant and transgenic complementation lines. (A) Representative images of four-weeks-old plants of the indicated genotypes grown in parallel on soil under short day (SD) conditions. Scale bar = 1 cm. (B) Immunoblot analysis of GFP-trap magnetic agarose beads (Chromotek) immunoprecipitated protein from *Np* driven complementation lines and total protein extracts from *35S* driven complementation lines, derived from plants of the indicated genotypes. Samples were separated on a 7.5 % SDS polyacryamide gel, blotted onto a nitrocellulose membrane and probed with α-GFP primary antibody and horseradish peroxidase coupled secondary antibody. Loading for total protein extracts was monitored by PonceauS (PonsS) staining of the membrane. (C) Plants of the indicated genotypes were infiltrated with a *Pst* DC3000 (ΔAvrPto/AvrPtoB) suspension of 1 x 10^5^ cfu ml^-1^. Colony-forming units (cfu) were quantified 1 hour (day 0) and three days after infiltration (day 3). Data is presented as boxplots, datapoints are colored based on replicates. Asterisks indicate statistically significant differences to Col-0 (Kurskal-Wallis test; Pairwise Wilcoxon tests with Benjamini-Hochberg correction, *P* < 0.05).

**Figure S3.**
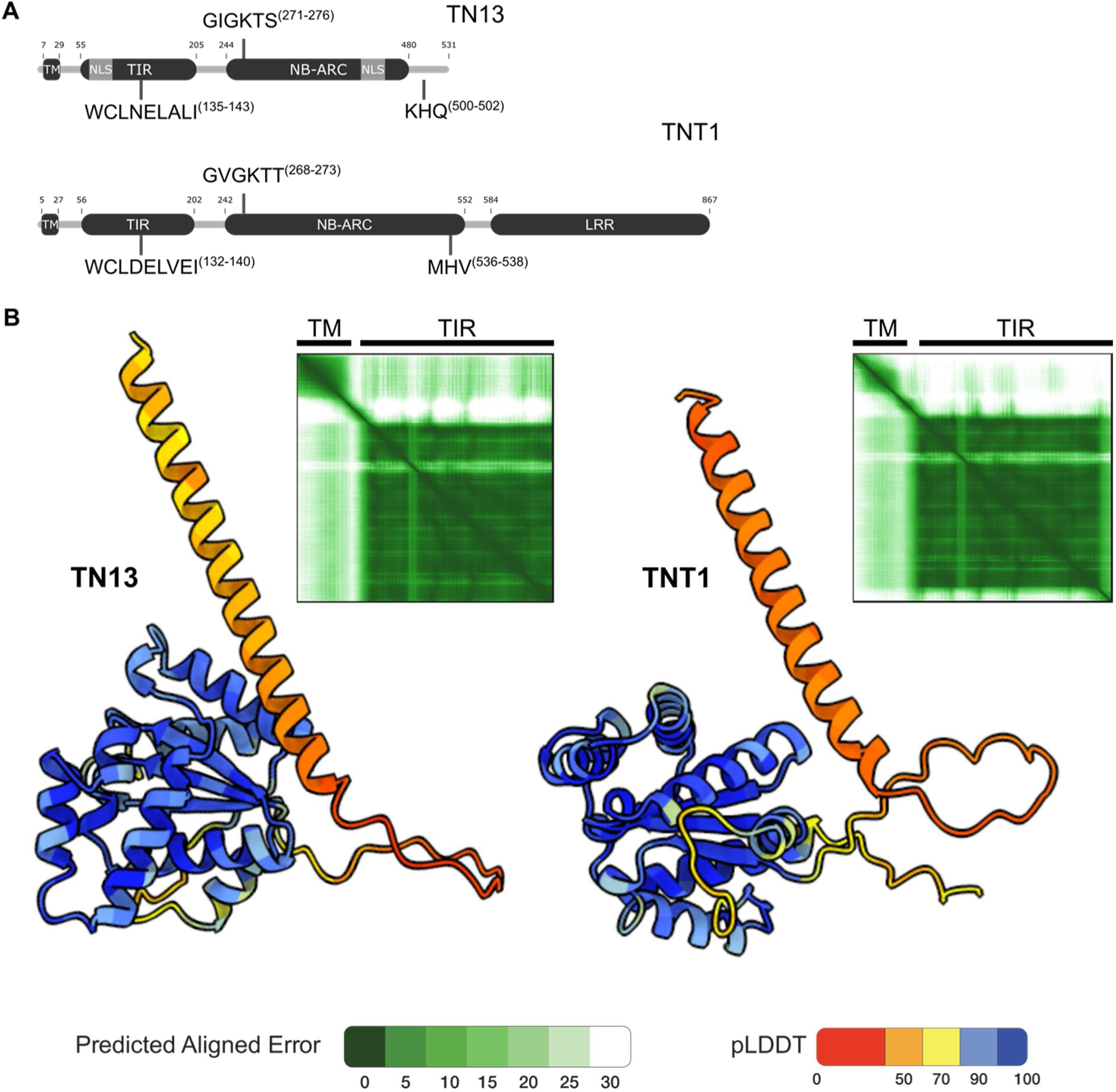
(A) Predicted domain structures and functional NLR motifs detected in TN13 and TNT1. The relative position of the TM, TIR and NB-ARC domains of TN13 and TNT1 are indicated. (B) Alphafold3 prediction of the N-terminal transmembrane (TM) and TIR domains of TN13 and TNT1. The predicted Local Distance Difference Test (pLDDT) as measure of per-residue local confidence, and the Predicted Aligned Error (PAE) to display the relative expected positonal error for each residue are shown.

**Figure S4.**
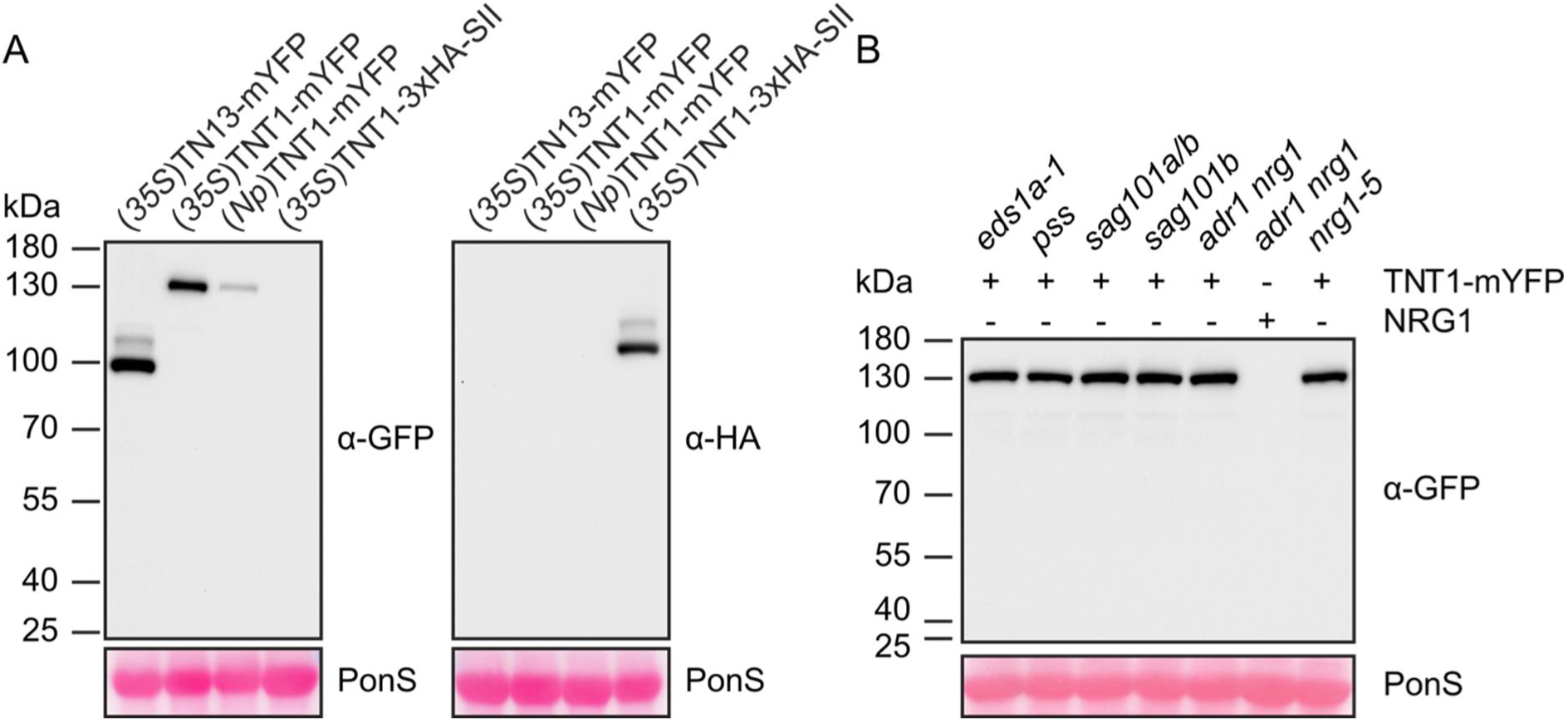
TNT1-mYFP protein accumulation upon *Agrobacterium*-mediated transient expression in *N. benthamiana eds1-1* mutant lines. (A) Immunoblot analysis of total protein extracts of *Nbeds1a-1* leave tissues, expressing TN13-mYFP, TNT1-mYFP or TNT1-3xHA-SII under control of the native (*Np*) or *35S* promoter. (B) Immunoblot analysis of total protein extracts from tissues of the indicated mutant lines, 5 days after infiltration of *Agrobacteria*. Samples were separated on a 7.5 % SDS polyacryamide gel, blotted onto nitrocellulose membrane and probed with α-GFP or α-HA primary antibody and horseradish peroxidase coupled secondary antibody. Loading was monitored by PonceauS (PonS) staining of the membrane.

**Figure S5.**
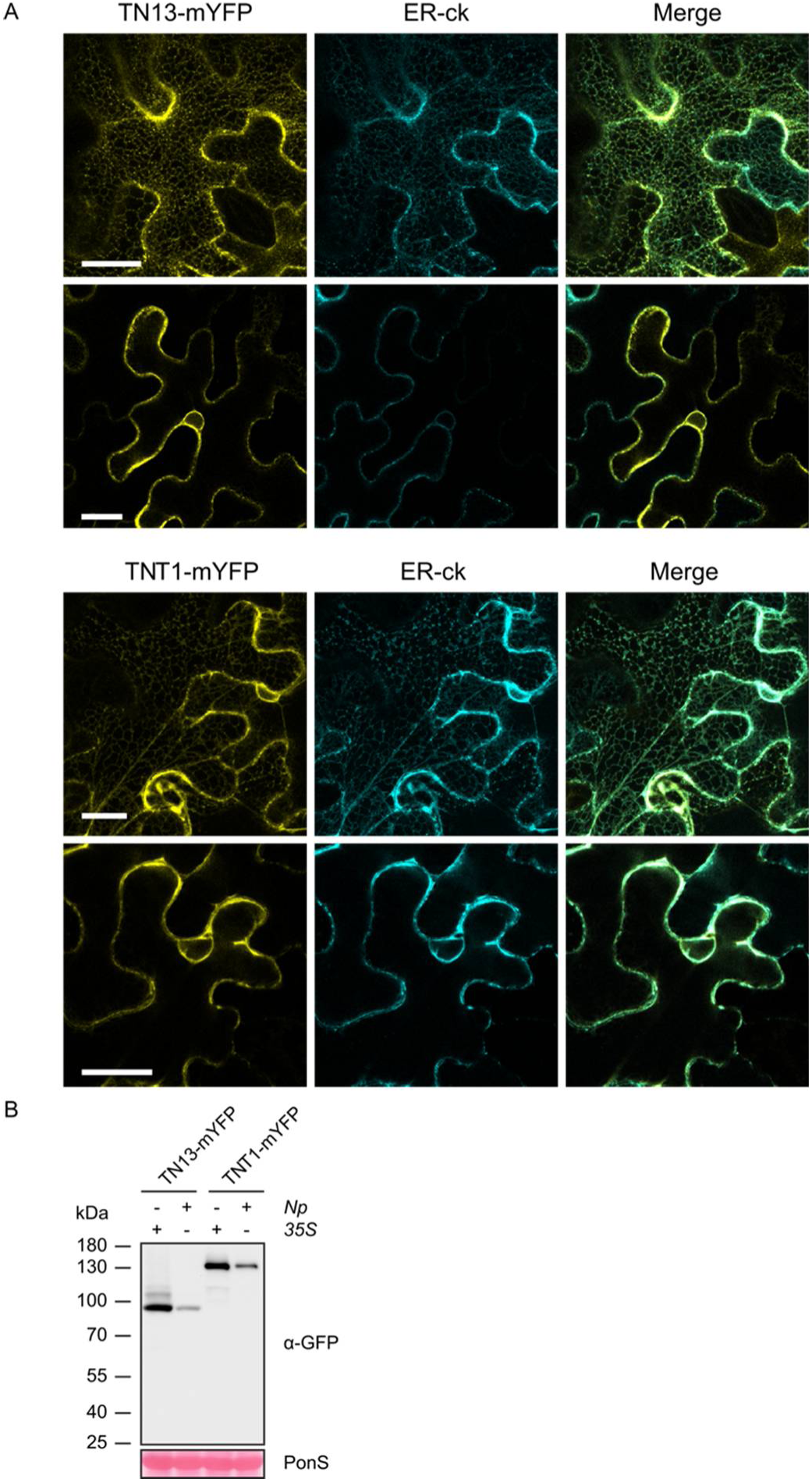
Localization of TN13-mYFP and TNT1-mYFP upon *Agrobacterium*-mediated transient expression in *Nbeds1a-1* leaves under control of the respective native promoters. (A) Confocal laser scanning microscopy was performed 2 days after *Agrobacterium* infiltration. The top panels show z-stacks, the panels below show a single confocal plane crossing the nucleus. TN13-mYFP and TNT1-mYFP are shown in yellow, the co-expressed CFP-tagged ER marker (ER-ck, Nelson *et al*., 2005) is shown in cyan. Scale bar = 25 µm. (B) Immunoblot analysis of total protein extracts derived from infiltrated *Nbeds1a-1* leave tissues expressing TN13-mYFP and TNT1-mYFP under control of the native (*Np*) or *35S* promoter. Samples were separated on a 7.5 % SDS polyacryamide gel, blotted onto a nitrocellulose membrane and probed with α-GFP primary antibody and horseradish peroxidase coupled secondary antibody. Loading was monitored by PonceauS (PonS) staining of the membrane.

**Figure S6.**
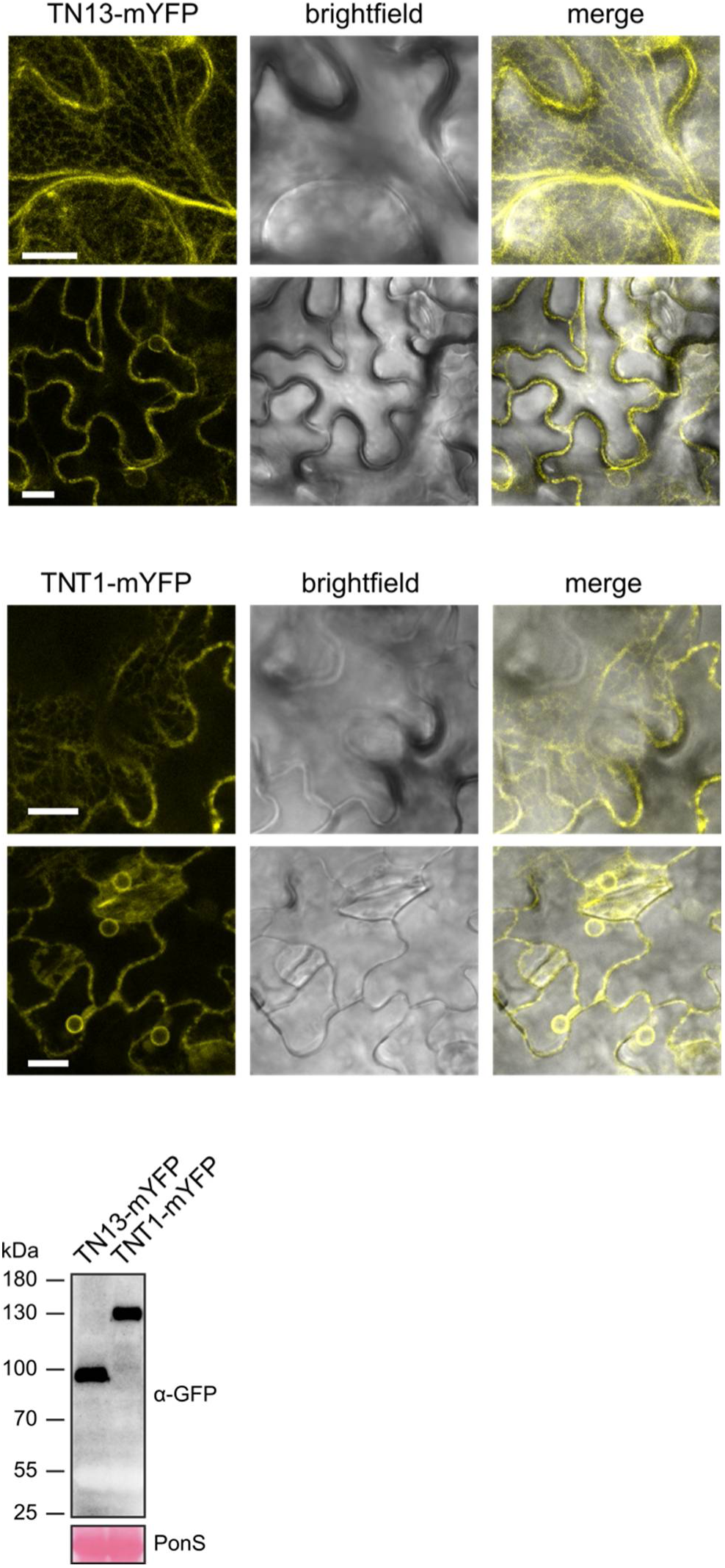
Localization of TN13-mYFP and TNT1-mYFP in stable transgenic *Arabidopsis* lines, expressing the fusion protein constructs under control of the *35S* promoter. Line #12-3 and #4-1 have been used respectively (see Figure S2). (A) Confocal laser scanning microscopy was performed on 2 weeks old plants, grown on ½ MS agar plates. The top panels show z-stacks, the panels below show a single confocal plane crossing nuclei. TN13-mYFP and TNT1-mYFP are shown in yellow. Scale bar = 25 µm. (B) Immunoblot analysis of total protein extracts, derived from plants used for CLSM in (A). Samples were separated on a 7.5 % SDS polyacryamide gel, blotted onto a nitrocellulose membrane and probed with α-GFP primary antibody and horseradish peroxidase coupled secondary antibody. Loading was monitored by PonceauS (PonS) staining of the membrane.

**Figure S7.**
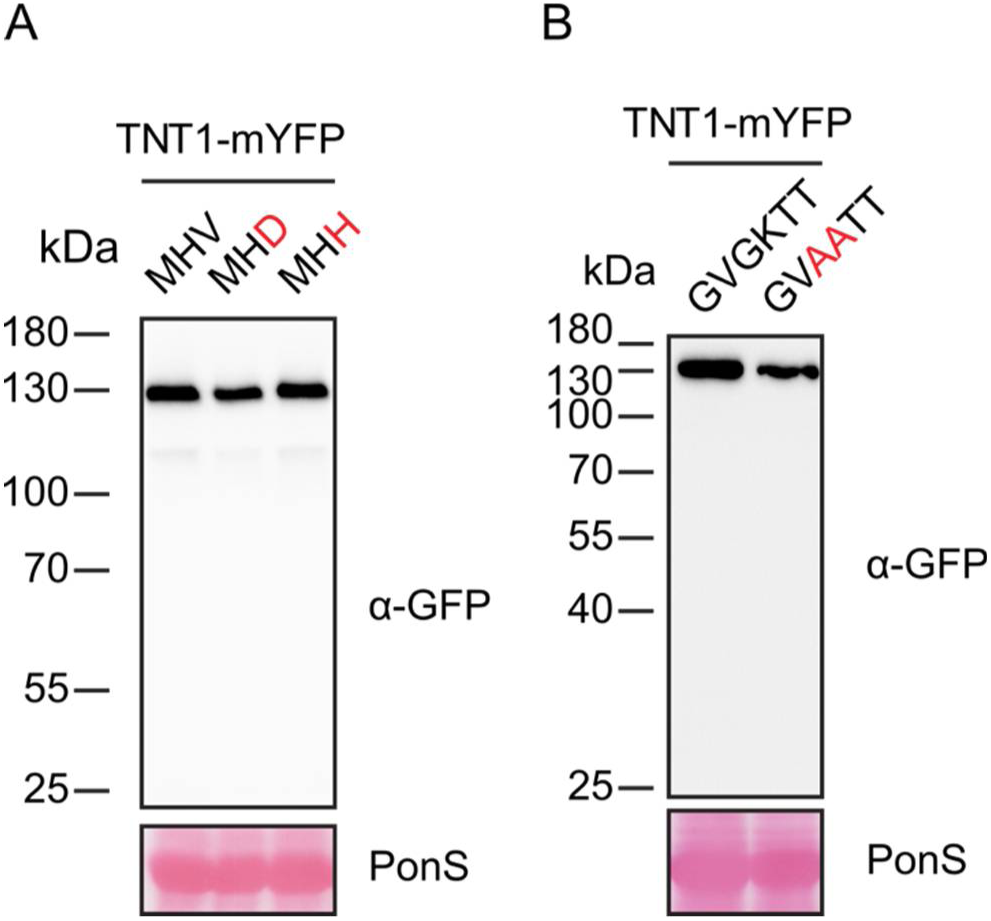
Immunoblot analysis of total protein extracts, derived from tissues transiently expressing TNT1-mYFP wild-type and mutant variants. Samples were taken 2 days after *Agrobacterium* infiltration of *N. benthamiana* wild-type plants for TNT1-mYFP mutant variants or *Nbeds1a-1* mutant plants for the TNT1-mYFP wild-type variant. All samples were separated on 7.5 % SDS polyacryamide gels, blotted onto nitrocellulose membranes and probed with the indicated primary antibodies. Horseradish peroxidase coupled secondary antibodies were used for detection. PonceauS (PonS) staining of the membrane was used as a loading control.

**Figure S8.**
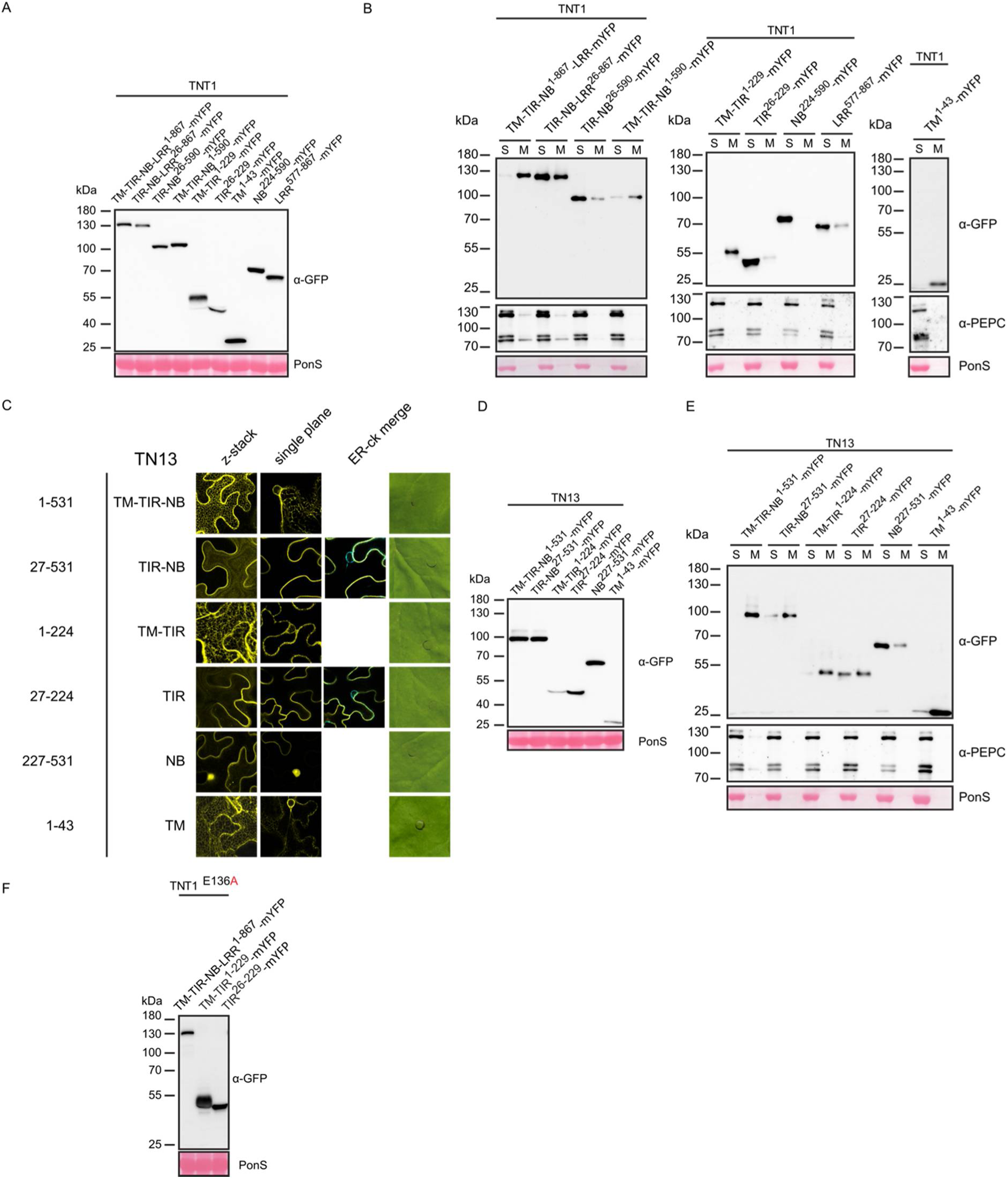
NLR domain truncation and mutation analysis. Immunoblot analysis of total protein extracts derived from tissues transiently expressing mYFP-tagged TNT1 wild-type and truncated versions (A), and TN13 wild-type and truncated versions (D). Samples were derived from *N. benthamiana* wild-type plants (for TN13 expression) or *Nbeds1a-1* mutant plants (for TNT1 expression) 2 days after infiltration of *Agrobacteria*. Immunoblots of protein extracts subjected to microsomal fractionation after expression of mYFP-tagged TNT1 wild-type and truncated versions (B) and TN13 wild-type and truncated versions (E). S = soluble fraction, M = membrane fraction. PEPC served as a cytoplasmic marker. (C) CLSM images of z-stacks and single confocal planes of the respective mYFP-tagged TN13 wild-type and truncated versions (shown in yellow), 2 days after infiltration of *Agrobacteria* into *N. benthamiana* wild-type plants. The ER marker fused to CFP (ER-ck, Nelson *et al*., 2005) was co-infiltrated to mark the nuclear envelope (shown in cyan). Confocal images showing the localization of the full length TN13 protein are also displayed in Figure 3A. For cell death assays, the infiltrated leave areas were photographed 5 days after infiltration of *Agrobacteria*. (F) Immunoblot analysis of total protein extracts, derived from infiltrated tissues expressing mYFP-tagged TNT1 full length or truncated versions, containing an E_136_A mutation of a predicted catalytic glutamate. Samples were taken 2 days after *Agrobacterium* infiltration of *N. benthamiana* wild-type plants. All protein samples were separated on 7.5 % SDS polyacrylamide gels, blotted onto nitrocellulose membranes and probed with α-GFP primary antibody and horseradish peroxidase coupled secondary antibody. Loading was monitored by PonceauS (PonS) staining of the membranes.

**Figure S9.**
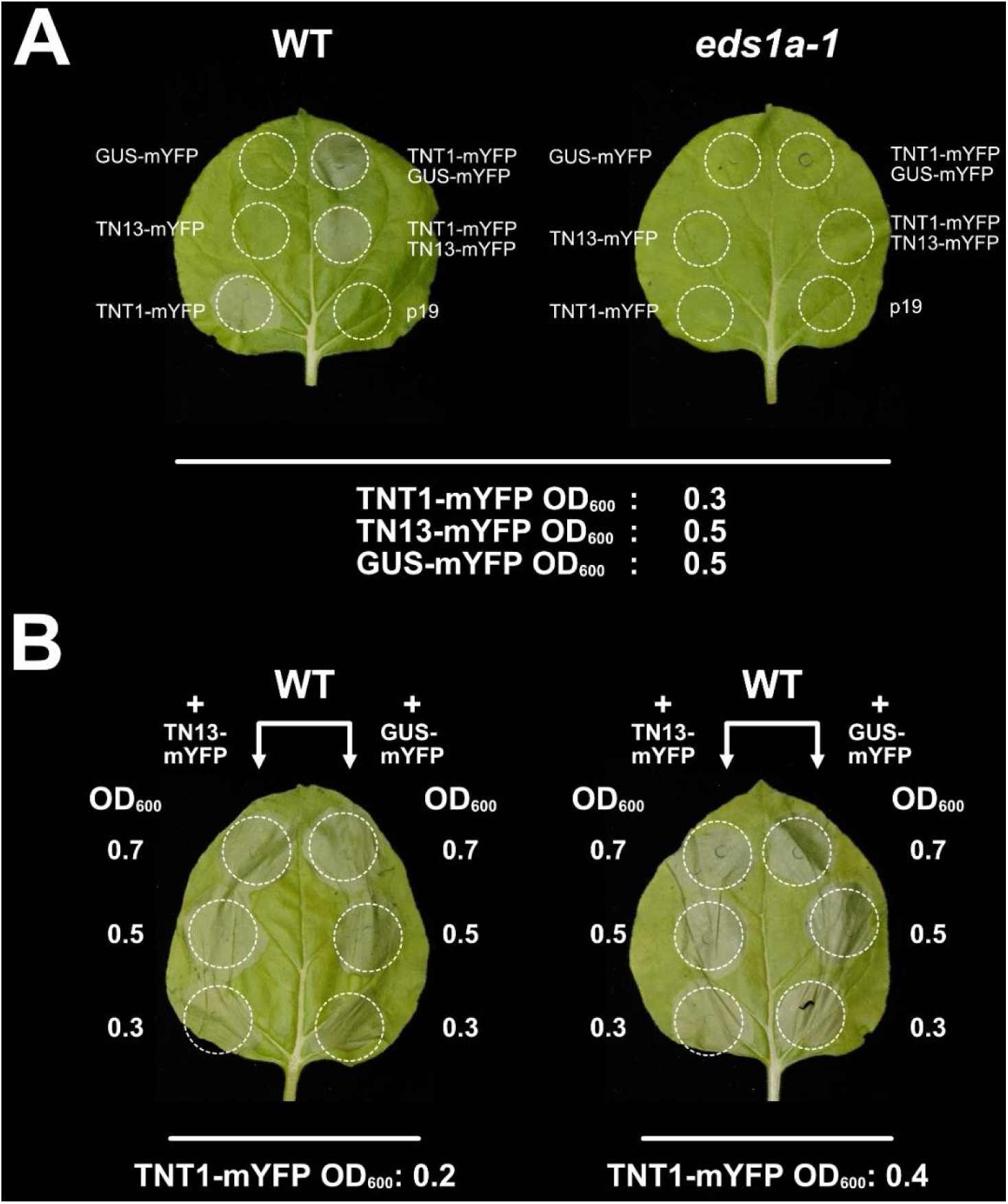
*Agrobacterium*-mediated transient co-expression of TNT1-mYFP and TN13-mYFP in wild-type *N. benthamiana* does not suppress TNT1 induced cell death. (A) Leaves of wild-type and *eds1a-1 N. benthamiana* plants were photographed 4 days after infiltration of *Agrobacterium* solutions of the indicated optical densities (OD_600_) for transient (co-)expression of mYFP tagged TN13, TNT1 or GUS under control of the *35S* promoter. The TNT1-induced cell death response depends on *NbEDS1*. Expression of GUS alone and of the silencing suppressor p19 alone served as additional controls. (B) Co-expression of TNT1-mYFP with either TN13-mYFP (left side of the leaf) or GUS-mYFP (right side of the leaf) in WT *N. benthamiana* leaves under control of the *35S* promoter. Leaves were photographed 4 days after infiltration of *Agrobacterium* solutions of the indicated optical densities (OD_600_). The silencing suppressor p19 was co-expressed with all combinations.

**Figure S10.**
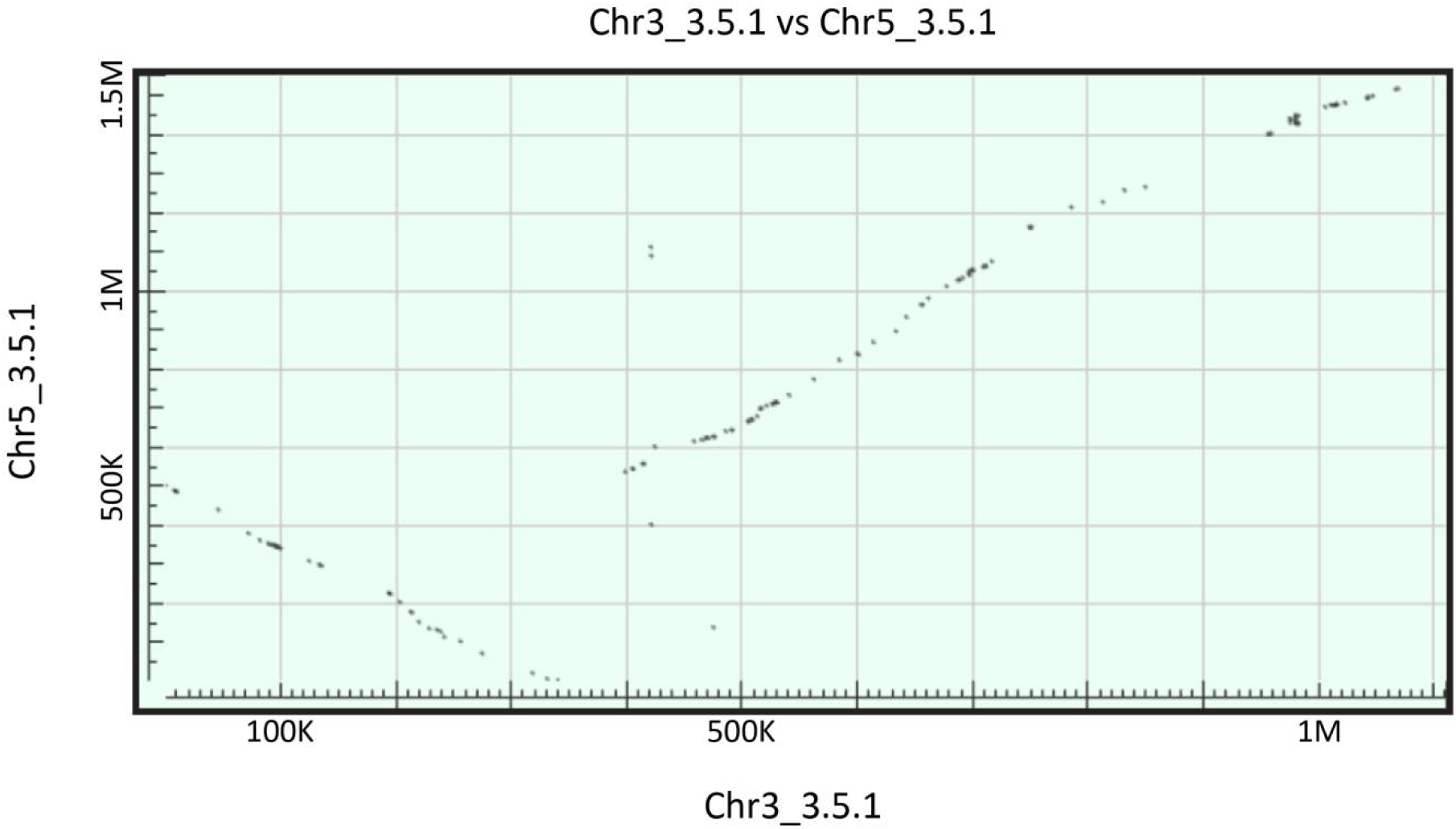
Dot plot matrix analysis of genomic regions. Dot matrix of alignment hits between the genomic sequences (based on TAIR10) of the regions between *AT3G01015* to *AT3G04350* on *Arabidopsis* chromosome 3 (Chr3_3.5.1), and *AT5G14060* to *AT5G18490* on chromosome 5 (Chr5_3.5.1), using the megablast settings in BLASTp. Numbers of bases are indicated.

**Figure S11.**
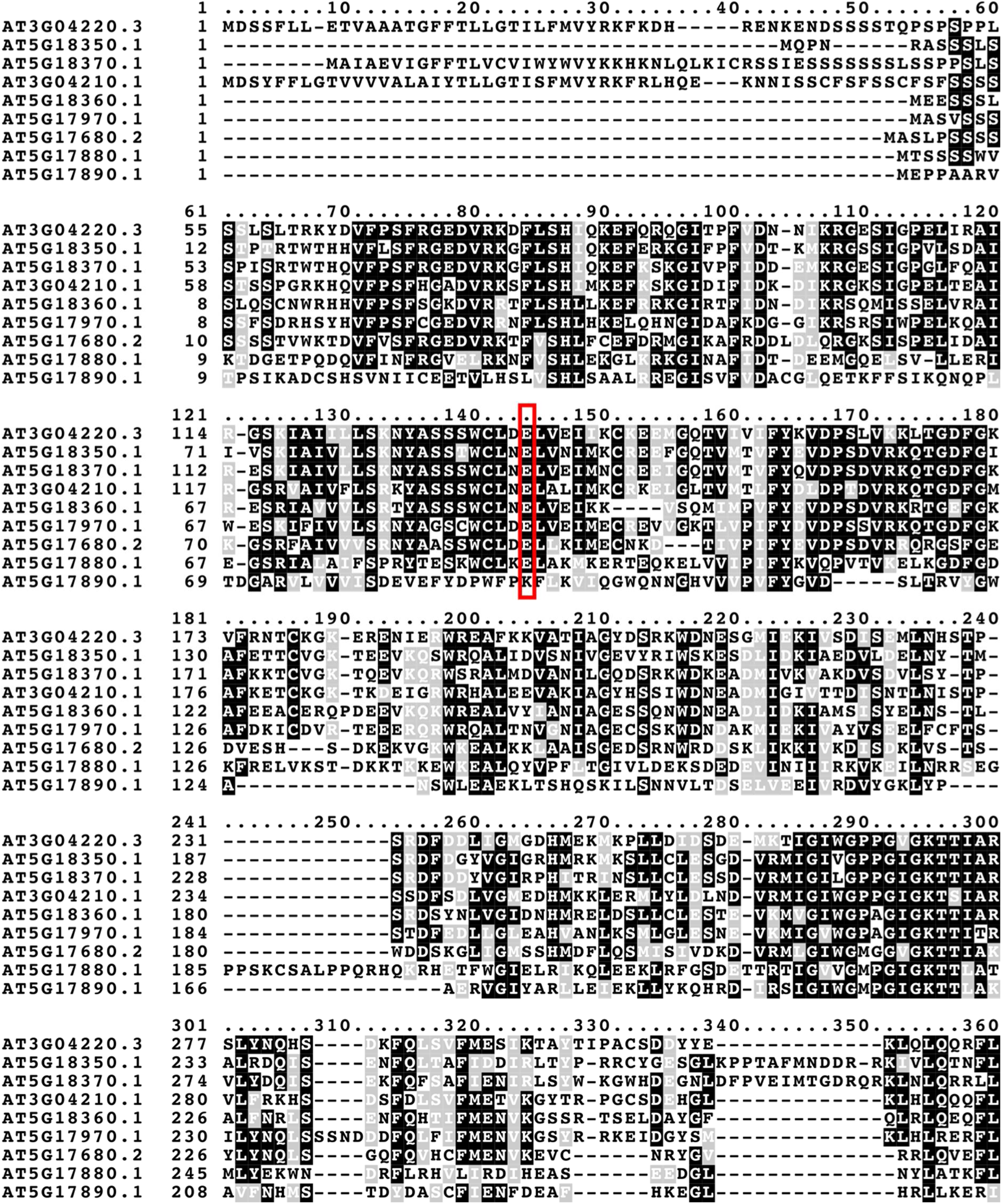
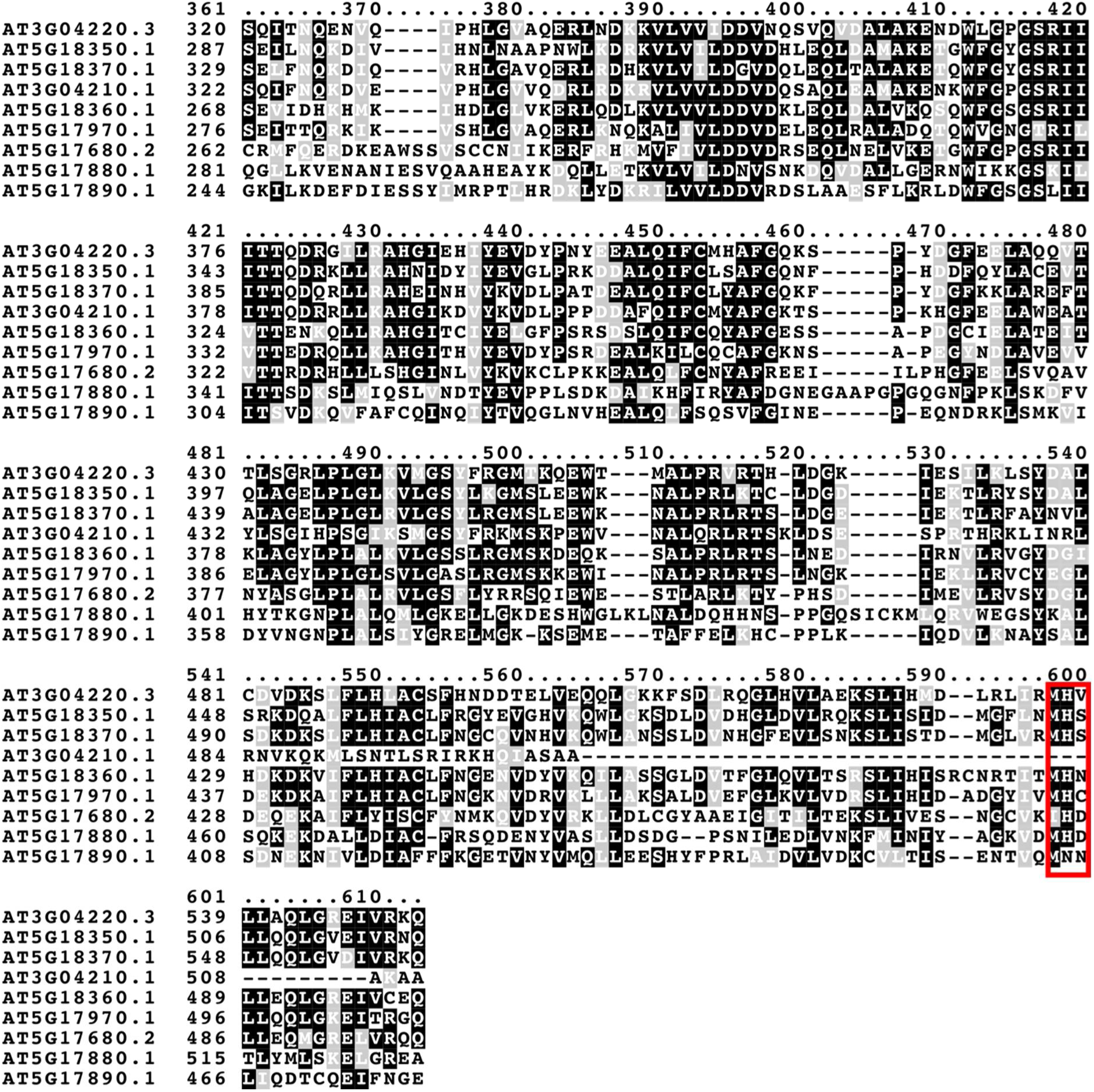
Protein sequence alignment of NLRs within the segmentally duplicated regions of chromosome 3 and 5. Alignment was performed using MAFFT, shading was performed using boxshade. Identical amino acids are highlighted in black and conserved amino acids in grey. The potential catalytic glutamate residue, and MHD-like motif are indicated by red boxes.

**Figure S12.**
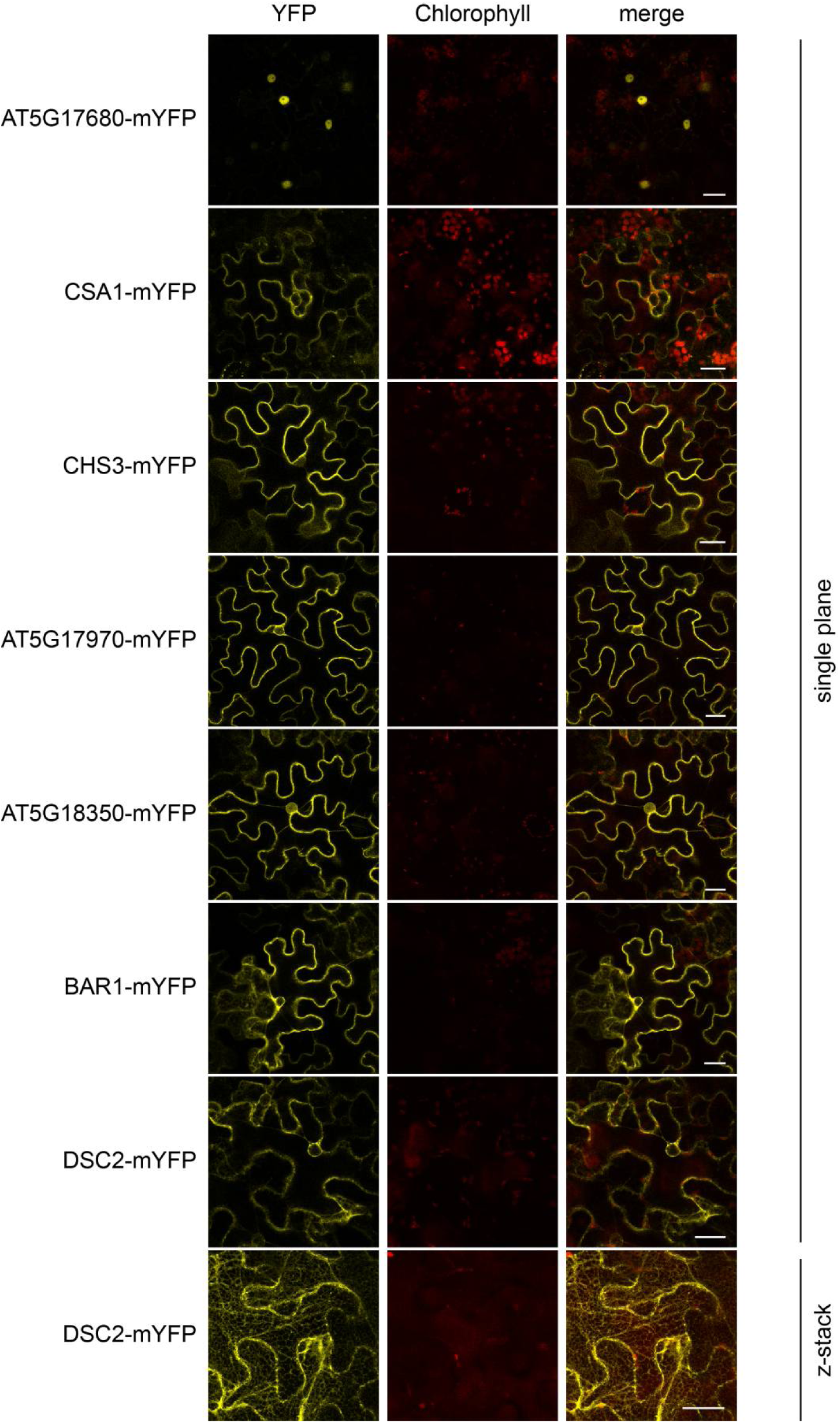
Localization of mYFP-tagged NLRs encoded in the segmentally duplicated cluster on chromosome 5. The localization of mYFP-tagged NLRs, encoded in the segmentally duplicated cluster on chromosome 5 was analized after *Agrobacterium*-mediated transient expression in *Nbeds1a-1* leaves, under control of the *35S* promoter. Confocal laser scanning microscopy was performed 2 days after infiltration of Agrobacteria. Z-stack and single confocal planes crossing the nucleus are shown. mYFP signals are shown in yellow, chlorophyll auto-fluorescence is shown in red. Scale bar = 25 µm.

**Figure S13.**
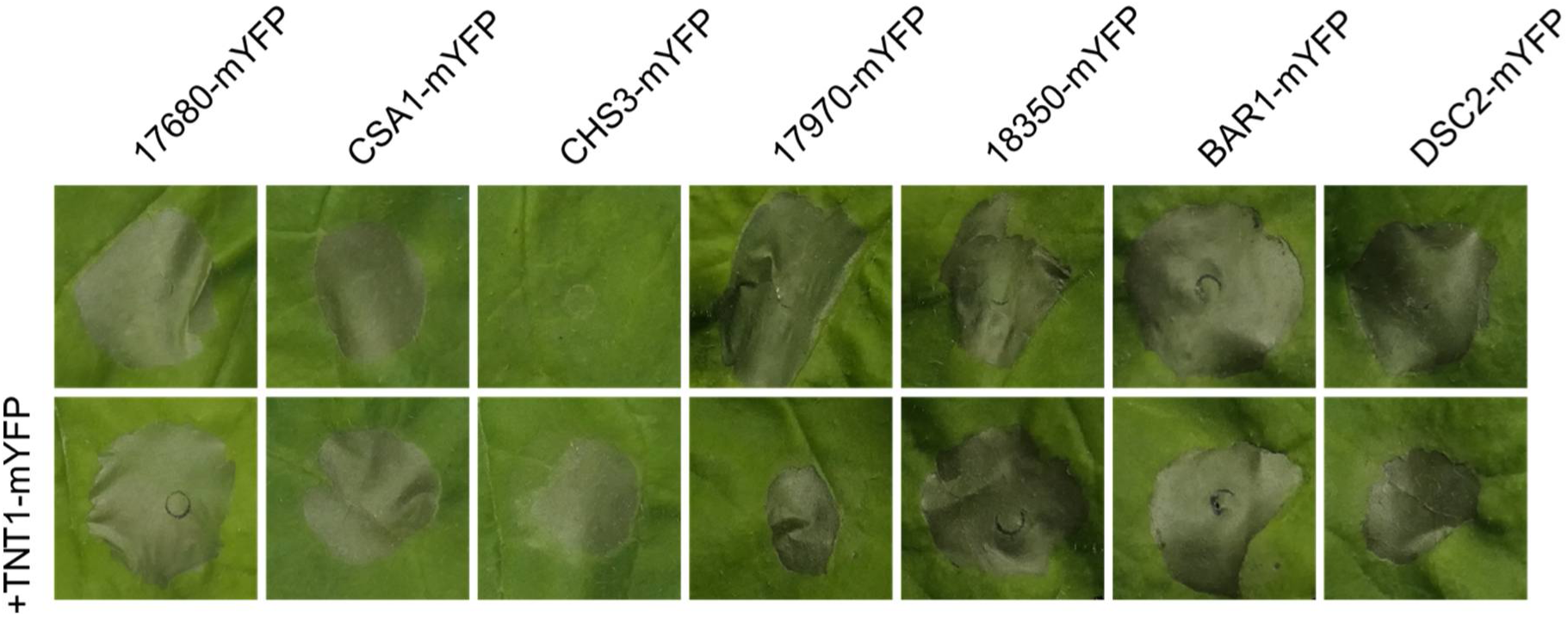
Cell death induction upon transient expression of NLRs encoded in the segmentally duplicated cluster on chromosome 5. *Agrobacterium*-mediated transient expression of mYFP-tagged NLRs, encoded in the segmentally duplicated cluster on chromosome 5 under control of the *35S* promoter. Upper panel shows individual expression of the NLRs and lower panel shows co-expression with TNT1-mYFP. Cell death was photographed 5 days after *Agrobacterium*-infiltration.

**Figure S14.**
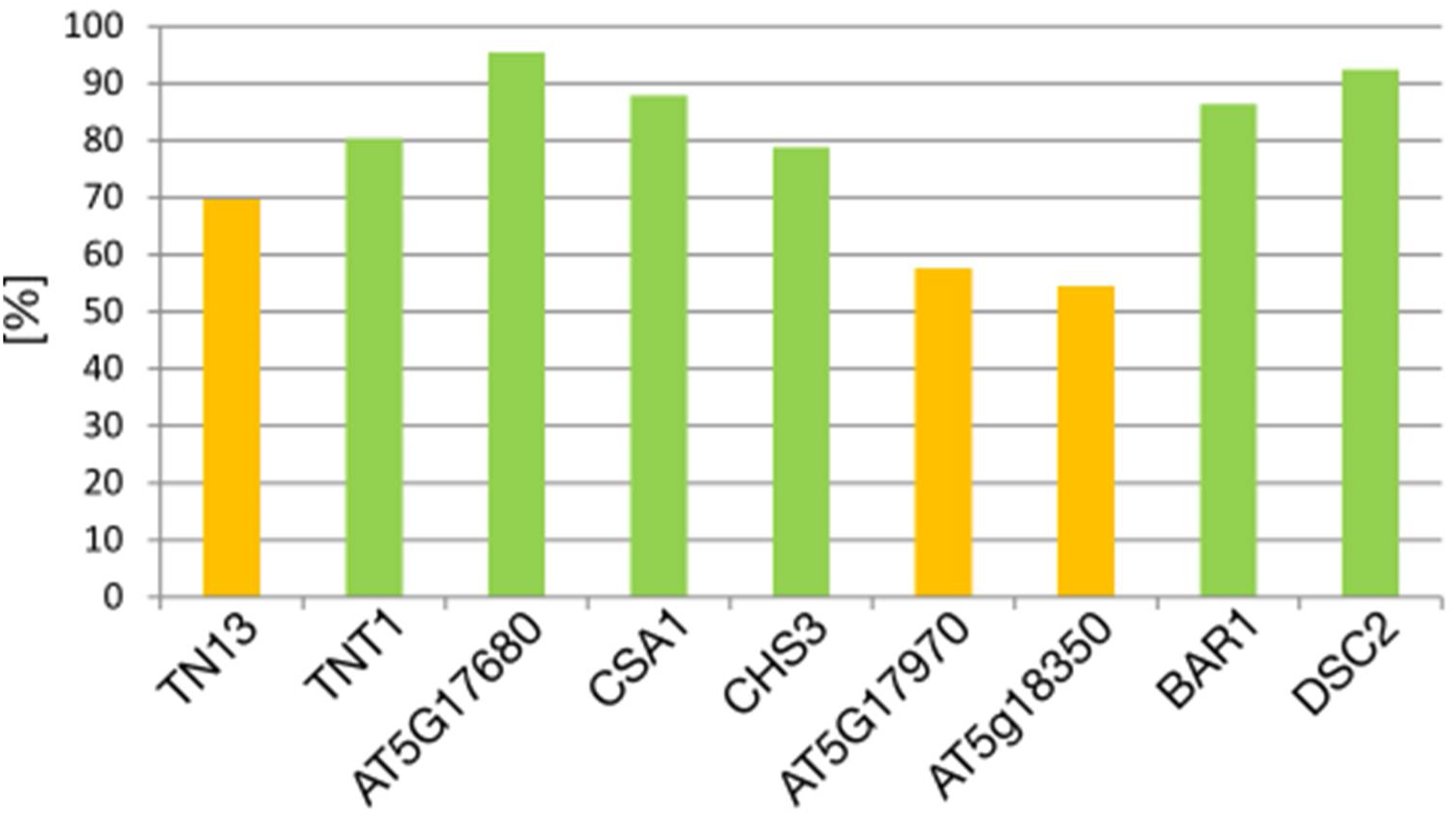
NLR presence analysis within *Arabidopsis* accessions. Presence (in %) of the respective orthogroups within the 65 *Arabidopsis* accessions used by Van de Weyer *et al*. (2019) to define core (>79 %, green), shell (>20 %, yellow) or cloud (<20 %, red; not applicable) NLRs within the pan-genome NLRome of *Arabidopsis thaliana*.

**Table S1.**
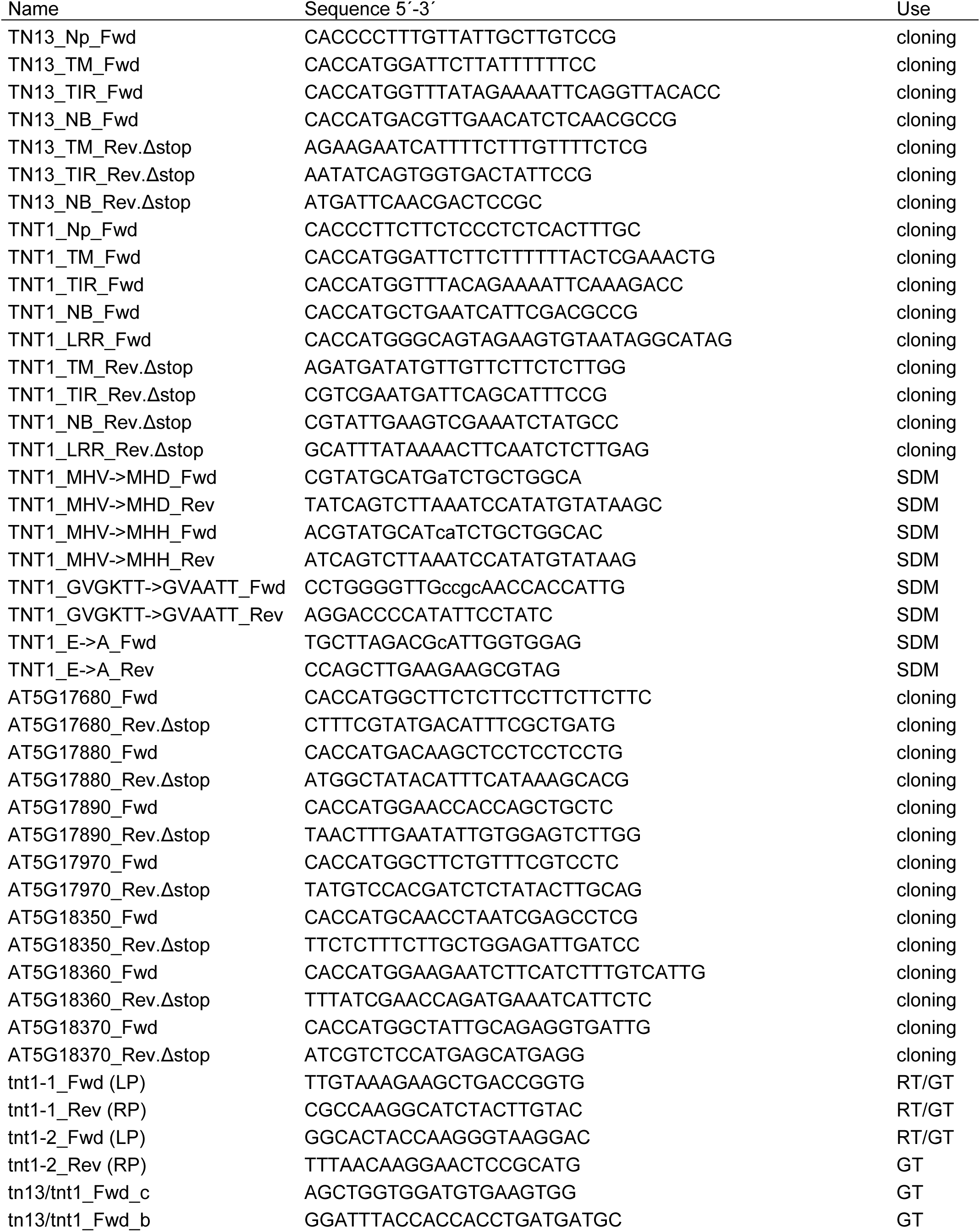
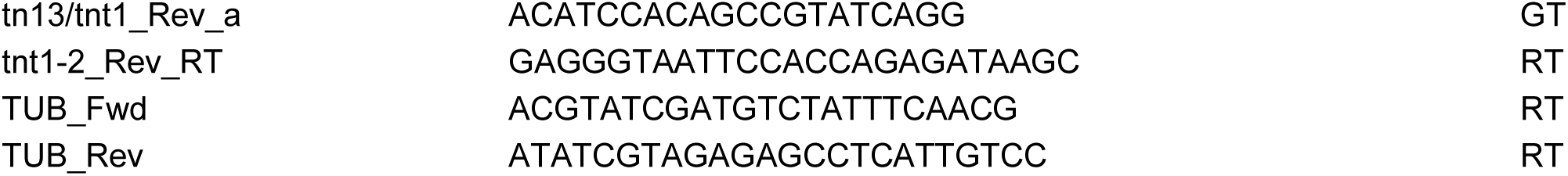
Primers used in this study. The name, sequence and use of each primer is indicated. SDM = site-directed mutagenesis, GT = genotyping, RT = RT-PCR.

## REFERENCES

1. Abramson, J., Adler, J., Dunger, J., et al. (2024) Accurate structure prediction of biomolecular interactions with AlphaFold 3. Nature, 630, 493–500.

2. Ahn, H.-K., Guo, G., Sklenar, J., et al. (2025) Recognition-dependent activation of the RRS1-R/RPS4 immune receptor complex. bioRxiv, 2025.04.11.646618. Available at: 10.1101/2025.04.11.646618v1.

3. Alcázar, R., Reth, M. von, Bautor, J., Chae, E., Weigel, D., Koornneef, M. and Parker, J.E. (2014) Analysis of a Plant Complex Resistance Gene Locus Underlying Immune-Related Hybrid Incompatibility and Its Occurrence in Nature. PLoS Genet., 10, e1004848.

4. Atanasov, K.E., Liu, C., Erban, A., Kopka, J., Parker, J.E. and Alcázar, R. (2018) NLR Mutations Suppressing Immune Hybrid Incompatibility and Their Effects on Disease Resistance. Plant Physiol., 177, 1152.

5. Barragan, A.C. and Weigel, D. (2021) Plant NLR diversity: the known unknowns of pan-NLRomes. Plant Cell, 33, 814–831.

6. Bartsch, M., Gobbato, E., Bednarek, P., Debey, S., Schultze, J.L., Bautor, J. and Parker, J.E. (2006) Salicylic Acid–Independent ENHANCED DISEASE SUSCEPTIBILITY1 Signaling in Arabidopsis Immunity and Cell Death Is Regulated by the Monooxygenase FMO1 and the Nudix Hydrolase NUDT7. Plant Cell, 18, 1038–1051.

7. Baumgarten, A., Cannon, S., Spangler, R. and May, G. (2003) Genome-Level Evolution of Resistance Genes in Arabidopsis thaliana. Genetics, 165, 309.

8. Bendahmane, A., Farnham, G., Moffett, P. and Baulcombe, D.C. (2002) Constitutive gain-of-function mutants in a nucleotide binding site–leucine rich repeat protein encoded at the Rx locus of potato. Plant J., 32, 195–204.

9. Bentham, A.R., Zdrzałek, R., De la Concepcion, J.C. and Banfield, M.J. (2018) Uncoiling CNLs: Structure/Function Approaches to Understanding CC Domain Function in Plant NLRs. Plant Cell Physiol., 59, 2398–2408.

10. Bernoux, M., Ve, T., Williams, S., et al. (2011) Structural and Functional Analysis of a Plant Resistance Protein TIR Domain Reveals Interfaces for Self-Association, Signaling, and Autoregulation. Cell Host Microbe, 9, 200–211.

11. Bomblies, K., Lempe, J., Epple, P., Warthmann, N., Lanz, C., Dangl, J.L. and Weigel, D. (2007) Autoimmune Response as a Mechanism for a Dobzhansky-Muller-Type Incompatibility Syndrome in Plants. PLoS Biol., 5, e236.

12. Bonardi, V., Tang, S., Stallmann, A., Roberts, M., Cherkis, K. and Dangl, J.L. (2011) Expanded functions for a family of plant intracellular immune receptors beyond specific recognition of pathogen effectors. Proceedings of the National Academy of Sciences, 108, 16463.

13. Cannon, S.B., Mitra, A., Baumgarten, A., Young, N.D. and May, G. (2004) The roles of segmental and tandem gene duplication in the evolution of large gene families in Arabidopsis thaliana. BMC Plant Biol., 4, 10.

14. Castel, B., Ngou, P.-M., Cevik, V., Redkar, A., Kim, D.-S., Yang, Y., Ding, P. and Jones, J.D.G. (2019) Diverse NLR immune receptors activate defence via the RPW8-NLR NRG1. New Phytol., 222, 966–980.

15. Césari, S., Kanzaki, H., Fujiwara, T., et al. (2014) The NB - LRR proteins RGA 4 and RGA 5 interact functionally and physically to confer disease resistance. EMBO J., 33, 1941–1959.

16. Chae, E., Bomblies, K., Kim, S.-T., et al. (2014) Species-wide Genetic Incompatibility Analysis Identifies Immune Genes as Hot Spots of Deleterious Epistasis. Cell, 159, 1341–1351.

17. Clough, S.J. and Bent, A.F. (1998) Floral dip: a simplified method for Agrobacterium-mediated transformation of Arabidopsis thaliana. Plant J., 16, 735–743.

18. Contreras, M.P., Lüdke, D., Pai, H., Toghani, A. and Kamoun, S. (2023) NLR receptors in plant immunity: making sense of the alphabet soup. EMBO Rep., 24, e57495.

19. Cui, H., Qiu, J., Zhou, Y., Bhandari, D.D., Zhao, C., Bautor, J. and Parker, J.E. (2018) Antagonism of Transcription Factor MYC2 by EDS1/PAD4 Complexes Bolsters Salicylic Acid Defense in Arabidopsis Effector-Triggered Immunity. Mol. Plant, 11, 1053–1066.

20. Dangl, J.L., Horvath, D.M. and Staskawicz, B.J. (2013) Pivoting the Plant Immune System from Dissection to Deployment. Science, 341, 746–751.

21. Danot, O., Marquenet, E., Vidal-Ingigliardi, D. and Richet, E. (2009) Wheel of Life, Wheel of Death: A Mechanistic Insight into Signaling by STAND Proteins. Structure, 17, 172–182.

22. Dong, O.X., Tong, M., Bonardi, V., El Kasmi, F., Woloshen, V., Wünsch, L.K., Dangl, J.L. and Li, X. (2016) TNL-mediated immunity in Arabidopsis requires complex regulation of the redundant ADR1 gene family. New Phytol., 210, 960– 973.

23. Feehan, J.M., Wang, J., Sun, X., Choi, J., Ahn, H.-K., Ngou, B.P.M., Parker, J.E. and Jones, J.D.G. (2023) Oligomerization of a plant helper NLR requires cell-surface and intracellular immune receptor activation. Proc. Natl. Acad. Sci. U. S. A., 120, e2210406120.

24. Feys, B.J., Wiermer, M., Bhat, R.A., Moisan, L.J., Medina-Escobar, N., Neu, C., Cabral, A. and Parker, J.E. (2005) Arabidopsis SENESCENCE-ASSOCIATED GENE101 Stabilizes and Signals within an ENHANCED DISEASE SUSCEPTIBILITY1 Complex in Plant Innate Immunity. The Plant Cell Online, 17, 2601–2613.

25. Gantner, J., Ordon, J., Kretschmer, C., Guerois, R. and Stuttmann, J. (2019) An EDS1-SAG101 Complex Is Essential for TNL-Mediated Immunity in Nicotiana benthamiana. Plant Cell, 31, 2456.

26. Genenncher, B., Wirthmueller, L., Roth, C., Klenke, M., Ma, L., Sharon, A. and Wiermer, M. (2016) Nucleoporin-Regulated MAP Kinase Signaling in Immunity to a Necrotrophic Fungal Pathogen. Plant Physiol., 172, 1293.

27. Horsefield, S., Burdett, H., Zhang, X., et al. (2019) NAD^+^ cleavage activity by animal and plant TIR domains in cell death pathways. Science, 365, 793.

28. Huang, S., Jia, A., Song, W., et al. Identification and receptor mechanism of TIR-catalyzed small molecules in plant immunity. Science, 377, eabq3297.

29. Huh, S.U., Cevik, V., Ding, P., Duxbury, Z., Ma, Y., Tomlinson, L., Sarris, P.F. and Jones, J.D.G. (2017) Protein-protein interactions in the RPS4/RRS1 immune receptor complex. PLoS Pathog., 13, e1006376.

30. Ibrahim, T., Yuen, E.L.H., Wang, H.-Y., et al. (2024) A helper NLR targets organellar membranes to trigger immunity. bioRxivorg, 2024.09.19.613839.

31. Jacob, P., Kim, N.H., Wu, F., et al. (2021) Plant “helper” immune receptors are Ca2+-permeable nonselective cation channels. Science, 373, 420–425.

32. Jia, A., Huang, S., Song, W., et al. (2022) TIR-catalyzed ADP-ribosylation reactions produce signaling molecules for plant immunity. Science, 377, eabq8180.

33. Jones, P., Binns, D., Chang, H.-Y., et al. (2014) InterProScan 5: genome-scale protein function classification. Bioinformatics, 30, 1236–1240.

34. Katoh, K., Misawa, K., Kuma, K. and Miyata, T. (2002) MAFFT: a novel method for rapid multiple sequence alignment based on fast Fourier transform. Nucleic Acids Res., 30, 3059–3066.

35. Katoh, K. and Standley, D.M. (2013) MAFFT Multiple Sequence Alignment Software Version 7: Improvements in Performance and Usability. Mol. Biol. Evol., 30, 772–780.

36. Kearse, M., Moir, R., Wilson, A., et al. (2012) Geneious Basic: An integrated and extendable desktop software platform for the organization and analysis of sequence data. Bioinformatics, 28, 1647–1649.

37. Kourelis, J. and Hoorn, R.A.L. van der (2018) Defended to the Nines: 25 Years of Resistance Gene Cloning Identifies Nine Mechanisms for R Protein Function. Plant Cell, 30, 285.

38. Kourelis, J., Sakai, T., Adachi, H. and Kamoun, S. (2021) RefPlantNLR is a comprehensive collection of experimentally validated plant disease resistance proteins from the NLR family. PLoS Biol., 19, e3001124.

39. Krogh, A., Larsson, B., Heijne, G. von and Sonnhammer, E.L.L. (2001) Predicting transmembrane protein topology with a hidden markov model: application to complete genomes11Edited by F. Cohen. J. Mol. Biol., 305, 567–580.

40. Laflamme, B., Dillon, M.M., Martel, A., Almeida, R.N.D., Desveaux, D. and Guttman, D.S. (2020) The pan-genome effector-triggered immunity landscape of a host-pathogen interaction. Science, 367, 763.

41. Lang, J., Genot, B., Bigeard, J. and Colcombet, J. (2022) MPK3 and MPK6 control salicylic acid signaling by up-regulating NLR receptors during pattern- and effector-triggered immunity. J. Exp. Bot., 73, 2190–2205.

42. Lapin, D., Kovacova, V., Sun, X., et al. (2019) A Coevolved EDS1-SAG101-NRG1 Module Mediates Cell Death Signaling by TIR-Domain Immune Receptors. Plant Cell, 31, 2430.

43. Le Roux, C., Huet, G., Jauneau, A., et al. (2016) A Receptor Pair with an Integrated Decoy Converts Pathogen Disabling of Transcription Factors to Immunity. Cell, 161, 1074–1088.

44. Lee, J., Lee, H., Kim, J., Lee, S., Kim, D.H., Kim, S. and Hwang, I. (2011) Both the Hydrophobicity and a Positively Charged Region Flanking the C-Terminal Region of the Transmembrane Domain of Signal-Anchored Proteins Play Critical Roles in Determining Their Targeting Specificity to the Endoplasmic Reticulum or Endosymbiotic Organelles in Arabidopsis Cells. Plant Cell, 23, 1588–1607.

45. Lee, R.R.Q. and Chae, E. (2020) Variation Patterns of NLR Clusters in Arabidopsis thaliana Genomes. Plant Commun, 1, 100089.

46. Leipe, D.D., Koonin, E.V. and Aravind, L. (2004) STAND, a Class of P-Loop NTPases Including Animal and Plant Regulators of Programmed Cell Death: Multiple, Complex Domain Architectures, Unusual Phyletic Patterns, and Evolution by Horizontal Gene Transfer. J. Mol. Biol., 343, 1–28.

47. Leister, D. (2004) Tandem and segmental gene duplication and recombination in the evolution of plant disease resistance genes. Trends Genet., 20, 116–122.

48. Li, X., Clarke, J., Zhang, Y. and Dong, X. (2001) Activation of an EDS1-Mediated R-Gene Pathway in the snc1 Mutant Leads to Constitutive, NPR1-Independent Pathogen Resistance. Mol. Plant. Microbe. Interact., 14, 1131–1139.

49. Liang, W., Wersch, S. van, Tong, M. and Li, X. (2019) TIR-NB-LRR immune receptor SOC3 pairs with truncated TIR-NB protein CHS1 or TN2 to monitor the homeostasis of E3 ligase SAUL1. New Phytol., 221, 2054–2066.

50. Lolle, S., Greeff, C., Petersen, K., et al. (2017) Matching NLR Immune Receptors to Autoimmunity in camta3 Mutants Using Antimorphic NLR Alleles. Cell Host Microbe, 21, 518–529.e4.

51. Lüdke, D., Yan, Q., Rohmann, P.F.W. and Wiermer, M. (2022) NLR we there yet? Nucleocytoplasmic coordination of NLR-mediated immunity. New Phytol., 236, 24–42.

52. Ma, Y., Guo, H., Hu, L., et al. (2018) Distinct modes of derepression of an Arabidopsis immune receptor complex by two different bacterial effectors. Proceedings of the National Academy of Sciences, 115, 10218.

53. Maruta, N., Burdett, H., Lim, B.Y.J., Hu, X., Desa, S., Manik, M.K. and Kobe, B. (2022) Structural basis of NLR activation and innate immune signalling in plants. Immunogenetics, 74, 5–26.

54. Meyers, B.C., Kozik, A., Griego, A., Kuang, H. and Michelmore, R.W. (2003) Genome-Wide Analysis of NBS-LRR–Encoding Genes in Arabidopsis. Plant Cell, 15, 809–834.

55. Meyers, B.C., Morgante, M. and Michelmore, R.W. (2002) TIR-X and TIR-NBS proteins: two new families related to disease resistance TIR-NBS-LRR proteins encoded in Arabidopsis and other plant genomes. Plant J., 32, 77–92.

56. Michael Weaver, L., Swiderski, M.R., Li, Y. and Jones, J.D.G. (2006) The Arabidopsis thaliana TIR-NB-LRR R-protein, RPP1A; protein localization and constitutive activation of defence by truncated alleles in tobacco and Arabidopsis. Plant J., 47, 829–840.

57. Nandety, R.S., Caplan, J.L., Cavanaugh, K., Perroud, B., Wroblewski, T., Michelmore, R.W. and Meyers, B.C. (2013) The Role of TIR-NBS and TIR-X Proteins in Plant Basal Defense Responses. Plant Physiol., 162, 1459–1472.

58. Nelson, B.K., Cai, X. and Nebenführ, A. (2007) A multicolored set of in vivo organelle markers for co-localization studies in Arabidopsis and other plants. Plant J., 51, 1126–1136.

59. Nishimura, M.T., Anderson, R.G., Cherkis, K.A., et al. (2017) TIR-only protein RBA1 recognizes a pathogen effector to regulate cell death in Arabidopsis. Proceedings of the National Academy of Sciences, 114, E2053.

60. Ntoukakis, V., Balmuth, A.L., Mucyn, T.S., Gutierrez, J.R., Jones, A.M.E. and Rathjen, J.P. (2013) The Tomato Prf Complex Is a Molecular Trap for Bacterial Effectors Based on Pto Transphosphorylation. PLoS Pathog., 9, e1003123.

61. Ordon, J., Gantner, J., Kemna, J., Schwalgun, L., Reschke, M., Streubel, J., Boch, J. and Stuttmann, J. (2017) Generation of chromosomal deletions in dicotyledonous plants employing a user-friendly genome editing toolkit. Plant J., 89, 155–168.

62. Peart, J.R., Mestre, P., Lu, R., Malcuit, I. and Baulcombe, D.C. (2005) NRG1, a CC-NB-LRR Protein, together with N, a TIR-NB-LRR Protein, Mediates Resistance against Tobacco Mosaic Virus. Curr. Biol., 15, 968–973.

63. Pettersen, E.F., Goddard, T.D., Huang, C.C., Meng, E.C., Couch, G.S., Croll, T.I., Morris, J.H. and Ferrin, T.E. (2021) UCSF ChimeraX: Structure visualization for researchers, educators, and developers. Protein Sci., 30, 70–82.

64. Price, M.N., Dehal, P.S. and Arkin, A.P. (2010) FastTree 2--approximately maximum-likelihood trees for large alignments. PLoS One, 5, e9490.

65. Qi, T., Seong, K., Thomazella, D.P.T., Kim, J.R., Pham, J., Seo, E., Cho, M.-J., Schultink, A. and Staskawicz, B.J. (2018) NRG1 functions downstream of EDS1 to regulate TIR-NLR-mediated plant immunity in Nicotiana benthamiana. Proceedings of the National Academy of Sciences, 115, E10979.

66. Reubold, T.F., Wohlgemuth, S. and Eschenburg, S. (2011) Crystal Structure of Full-Length Apaf-1: How the Death Signal Is Relayed in the Mitochondrial Pathway of Apoptosis. Structure, 19, 1074–1083.

67. Richly, E., Kurth, J. and Leister, D. (2002) Mode of Amplification and Reorganization of Resistance Genes During Recent Arabidopsis thaliana Evolution. Mol. Biol. Evol., 19, 76–84.

68. Roth, C., Lüdke, D., Klenke, M., Quathamer, A., Valerius, O., Braus, G.H. and Wiermer, M. (2017) The truncated NLR protein TIR-NBS13 is a MOS6/IMPORTIN-α3 interaction partner required for plant immunity. Plant J., 92, 808–821.

69. Saile, S.C., Jacob, P., Castel, B., Jubic, L.M., Salas-Gonzáles, I., Bäcker, M., Jones, J.D.G., Dangl, J.L. and El Kasmi, F. (2020) Two unequally redundant “helper” immune receptor families mediate Arabidopsis thaliana intracellular “sensor” immune receptor functions. PLoS Biol., 18, e3000783.

70. Sarris, P.F., Duxbury, Z., Huh, S.U., et al. (2015) A Plant Immune Receptor Detects Pathogen Effectors that Target WRKY Transcription Factors. Cell, 161, 1089– 1100.

71. Schindelin, J., Arganda-Carreras, I., Frise, E., et al. (2012) Fiji: an open-source platform for biological-image analysis. Nat. Methods, 9, 676–682.

72. Schreiber, K.J., Bentham, A., Williams, S.J., Kobe, B. and Staskawicz, B.J. (2016) Multiple Domain Associations within the Arabidopsis Immune Receptor RPP1 Regulate the Activation of Programmed Cell Death. PLoS Pathog., 12, e1005769.

73. Schulze, S., Yu, L., Hua, C., et al. (2022) The Arabidopsis TIR-NBS-LRR protein CSA1 guards BAK1-BIR3 homeostasis and mediates convergence of pattern- and effector-induced immune responses. Cell Host Microbe, 30, 1717–1731.e6.

74. Selvaraj, M., Toghani, A., Pai, H., et al. (2024) Activation of plant immunity through conversion of a helper NLR homodimer into a resistosome. PLoS Biol., 22, e3002868.

75. Shao, Z.-Q., Xue, J.-Y., Wu, P., Zhang, Y.-M., Wu, Y., Hang, Y.-Y., Wang, B. and Chen, J.-Q. (2016) Large-Scale Analyses of Angiosperm Nucleotide-Binding Site-Leucine-Rich Repeat Genes Reveal Three Anciently Diverged Classes with Distinct Evolutionary Patterns. Plant Physiol., 170, 2095.

76. Simillion, C., Vandepoele, K., Van Montagu, M.C.E., Zabeau, M. and Van de Peer, Y. (2002) The hidden duplication past of Arabidopsis thaliana. Proceedings of the National Academy of Sciences, 99, 13627.

77. Slootweg, E.J., Spiridon, L.N., Roosien, J., et al. (2013) Structural Determinants at the Interface of the ARC2 and Leucine-Rich Repeat Domains Control the Activation of the Plant Immune Receptors Rx1 and Gpa2. Plant Physiol., 162, 1510.

78. Staal, J., Kaliff, M., Dewaele, E., Persson, M. and Dixelius, C. (2008) RLM3, a TIR domain encoding gene involved in broad-range immunity of Arabidopsis to necrotrophic fungal pathogens. Plant J., 55, 188–200.

79. Stuttmann, J., Peine, N., Garcia, A.V., et al. (2016) Arabidopsis thaliana DM2h (R8) within the Landsberg RPP1-like Resistance Locus Underlies Three Different Cases of EDS1-Conditioned Autoimmunity. PLoS Genet., 12, e1005990.

80. Sukarta, O.C.A., Slootweg, E.J. and Goverse, A. (2016) Structure-informed insights for NLR functioning in plant immunity. X chromosome inactivation, 56, 134– 149.

81. Takken, F.L.W., Albrecht, M. and Tameling, W.I.L. (2006) Resistance proteins: molecular switches of plant defence. Biotic interactions / edited by Anne Osbourn and Sheng Yang He, 9, 383–390.

82. Tameling, W.I.L., Vossen, J.H., Albrecht, M., Lengauer, T., Berden, J.A., Haring, M.A., Cornelissen, B.J.C. and Takken, F.L.W. (2006) Mutations in the NB-ARC Domain of I-2 That Impair ATP Hydrolysis Cause Autoactivation. Plant Physiol., 140, 1233.

83. Thomma, B.P.H.J., Nürnberger, T. and Joosten, M.H.A.J. (2011) Of PAMPs and Effectors: The Blurred PTI-ETI Dichotomy. Plant Cell, 23, 4–15.

84. Tian, H., Wu, Z., Chen, S., et al. (2021) Activation of TIR signalling boosts pattern-triggered immunity. Nature, 598, 500–503.

85. Toghani, A. and Kamoun, S. (2024) Functional annotation of 180 RefSeq reference plant proteomes reveals a dataset of 113,684 NLR proteins. Available at: 10.5281/zenodo.13627395.

86. Tong, M., Kotur, T., Liang, W., et al. (2017) E3 ligase SAUL1 serves as a positive regulator of PAMP-triggered immunity and its homeostasis is monitored by immune receptor SOC3. New Phytol., 215, 1516–1532.

87. Van de Weyer, A.-L., Monteiro, F., Furzer, O.J., et al. (2019) A Species-Wide Inventory of NLR Genes and Alleles in Arabidopsis thaliana. Cell, 178, 1260–1272.e14.

88. Ve, T., Williams, S.J. and Kobe, B. (2015) Structure and function of Toll/interleukin-1 receptor/resistance protein (TIR) domains. Apoptosis, 20, 250–261.

89. Wagner, S., Stuttmann, J., Rietz, S., Guerois, R., Brunstein, E., Bautor, J., Niefind, K. and Parker, J.E. (2013) Structural Basis for Signaling by Exclusive EDS1 Heteromeric Complexes with SAG101 or PAD4 in Plant Innate Immunity. Cell Host Microbe, 14, 619–630.

90. Wan, L., Essuman, K., Anderson, R.G., et al. (2019) TIR domains of plant immune receptors are NAD^+^-cleaving enzymes that promote cell death. Science, 365, 799.

91. Wang, Jizong, Hu, M., Wang, Jia, et al. (2019) Reconstitution and structure of a plant NLR resistosome conferring immunity. Science, 364, eaav5870.

92. Wang, Jizong, Wang, Jia, Hu, M., et al. (2019) Ligand-triggered allosteric ADP release primes a plant NLR complex. Science, 364. Available at: 10.1126/science.aav5868.

93. Wang, W., Liu, N., Gao, C., Rui, L. and Tang, D. (2019) The Pseudomonas Syringae Effector AvrPtoB Associates With and Ubiquitinates Arabidopsis Exocyst Subunit EXO70B1. Front. Plant Sci., 10, 1027.

94. Wersch, S. van and Li, X. (2019) Stronger When Together: Clustering of Plant NLR Disease resistance Genes. Trends Plant Sci., 24, 688–699.

95. Wiermer, M., Feys, B.J. and Parker, J.E. (2005) Plant immunity: the EDS1 regulatory node. Biotic interactions, 8, 383–389.

96. Williams, S.J., Sohn, K.H., Wan, L., et al. (2014) Structural Basis for Assembly and Function of a Heterodimeric Plant Immune Receptor. Science, 344, 299–303.

97. Williams, S.J., Sornaraj, P., Ireland, E. deCourcy-, Menz, R.I., Kobe, B., Ellis, J.G., Dodds, P.N. and Anderson, P.A. (2011) An Autoactive Mutant of the M Flax Rust Resistance Protein Has a Preference for Binding ATP, Whereas Wild-Type M Protein Binds ADP. Molecular Plant-Microbe Interactions®, 24, 897– 906.

98. Wu, C.-H., Abd-El-Haliem, A., Bozkurt, T.O., Belhaj, K., Terauchi, R., Vossen, J.H. and Kamoun, S. (2017) NLR network mediates immunity to diverse plant pathogens. Proceedings of the National Academy of Sciences, 114, 8113.

99. Wu, C.-H. and Derevnina, L. (2023) The battle within: How pathogen effectors suppress NLR-mediated immunity. Curr. Opin. Plant Biol., 74, 102396.

100. Wu, Y., Xu, W., Zhao, G., et al. (2024) A canonical protein complex controls immune homeostasis and multipathogen resistance. Science, 386, 1405–1412.

101. Wu, Z., Li, M., Dong, O.X., Xia, S., Liang, W., Bao, Y., Wasteneys, G. and Li, X. (2019) Differential regulation of TNL-mediated immune signaling by redundant helper CNLs. New Phytol., 222, 938–953.

102. Xiao, S., Calis, O., Patrick, E., Zhang, G., Charoenwattana, P., Muskett, P., Parker, J.E. and Turner, J.G. (2005) The atypical resistance gene, RPW8, recruits components of basal defence for powdery mildew resistance in Arabidopsis. Plant J., 42, 95–110.

103. Yang, Y., Furzer, O.J., Fensterle, E.P., Lin, S., Zheng, Z., Kim, N.H., Wan, L. and Dangl, J.L. (2024) Paired plant immune CHS3-CSA1 receptor alleles form distinct hetero-oligomeric complexes. Science, 383, eadk3468.

104. Yang, Y., Kim, N.H., Cevik, V., Jacob, P., Wan, L., Furzer, O.J. and Dangl, J.L. (2022) Allelic variation in the Arabidopsis TNL CHS3/CSA1 immune receptor pair reveals two functional cell-death regulatory modes. Cell Host Microbe, 30, 1701–1716.e5.

105. Yu, D., Song, W., Tan, E.Y.J., et al. (2022) TIR domains of plant immune receptors are 2′,3′-cAMP/cGMP synthetases mediating cell death. Cell. Available at: https://www.sciencedirect.com/science/article/pii/S009286742200530X.

106. Yu, H., Xu, W., Chen, S., et al. (2024) Activation of a helper NLR by plant and bacterial TIR immune signaling. Science, 386, 1413–1420.

107. Zhang, Y., Wang, Y., Liu, J., Ding, Y., Wang, S., Zhang, X., Liu, Y. and Yang, S. (2017) Temperature-dependent autoimmunity mediated by chs1 requires its neighboring TNL gene SOC3. New Phytol., 213, 1330–1345.

108. Zhao, T., Rui, L., Li, J., et al. (2015) A Truncated NLR Protein, TIR-NBS2, Is Required for Activated Defense Responses in the exo70B1 Mutant. PLoS Genet., 11, e1004945.

109. Zhou, M., Li, Y., Hu, Q., Bai, X.-C., Huang, W., Yan, C., Scheres, S.H.W. and Shi, Y. (2015) Atomic structure of the apoptosome: mechanism of cytochrome c- and dATP-mediated activation of Apaf-1. Genes Dev., 29, 2349–2361.

